# Natural Cystatin C fragments inhibit GPR15-mediated HIV and SIV infection without interfering with GPR15L signaling

**DOI:** 10.1101/2020.10.26.355172

**Authors:** Manuel Hayn, Andrea Blötz, Armando Rodríguez, Solange Vidal, Nico Preising, Ludger Ständker, Sebastian Wiese, Christina M. Stürzel, Mirja Harms, Rüdiger Groß, Christoph Jung, Miriam Kiene, Beatrice H. Hahn, Timo Jacob, Stefan Pöhlmann, Wolf-Georg Forssmann, Jan Münch, Konstantin M. J. Sparrer, Klaus Seuwen, Frank Kirchhoff

## Abstract

GPR15 is a G protein-coupled receptor proposed to play a role in mucosal immunity that also serves as entry cofactor for HIV and SIV. To discover novel endogenous GPR15 ligands, we screened a hemofiltrate-derived peptide library for inhibitors of GPR15-mediated SIV infection. Our approach identified a C-terminal fragment of Cystatin C (CysC95-146) that specifically inhibits GPR15-dependent HIV-1, HIV-2 and SIV infection. In contrast, GPR15L, the chemokine ligand of GPR15, failed to inhibit virus infection. We found that Cystatin C fragments preventing GPR15-mediated viral entry do not interfere with GPR15L signaling and are generated by proteases activated at sites of inflammation. The antiretroviral activity of CysC95-146 was confirmed in primary CD4+ T cells and is conserved in simian hosts of SIV infection. Thus, we identified a potent endogenous inhibitor of GPR15-mediated HIV and SIV infection that does not interfere with the physiological function of this G protein-coupled receptor.

## INTRODUCTION

G protein-coupled receptors (GPCRs) constitute the largest family of membrane proteins involved in the transduction of signals from the extracellular environment into the cell and play key roles in immune responses, homeostasis, metabolism and organogenesis (Heng et al., 2013; Venkatakrishnan et al., 2013). Besides their physiological roles, some GPCRs also represent important coreceptors for HIV and/or SIV entry. HIV-1, the main causative agent of AIDS, utilizes C-C chemokine receptor 5 (CCR5) and CXC chemokine receptor 4 (CXCR4) as major entry cofactors (Alkhatib et al., 1996; Choe et al., 1996; Deng et al., 1996; Doranz et al., 1996; Dragic et al., 1996; Feng et al., 1996). The chemokine ligands CCL5 (also known as RANTES) and CXCL12 (also named SDF-1) inhibit CCR5-or CXCR4-mediated HIV-1 infection, respectively. Thus, we have previously taken advantage of HIV-1 entry to examine complex blood-derived peptide libraries for novel naturally occurring ligands of CCR5 and CXCR4 (Münch et al., 2014; Bosso et al., 2017). Initially, we identified a truncated form of the Chemokine C-C ligand 14 (CCL14) as a novel CCR5 agonist and potent inhibitor of CCR5-tropic HIV-1 strains (Detheux et al., 2000; Münch et al., 2002). More recently, we discovered a small fragment of human serum albumin (EPI-X4) as an effective and highly specific CXCR4 antagonist and inhibitor of CXCR4-tropic HIV-1 strains (Zirafi et al., 2015).

While HIV-1 coreceptor utilization is mainly restricted to CCR5 and CXCR4, HIV-2 and SIVs are more promiscuous in their entry cofactor usage. For example, many HIV-2 and SIV strains utilize BOB/GPR15 and Bonzo/STRL-33/CXCR6 in addition to CCR5 and/or CXCR4 for viral entry into CD4^+^ target cells (Deng et al., 1997; Farzan et al., 1997; Mörner et al., 1999; Owen et al., 1998; Pöhlmann et al., 2000; Riddick et al., 2010, 2015). GPR15 is a GPCR reported to regulate T cell trafficking to the colon that may play a role in intestinal homeostasis and inflammation (Kim et al., 2013; Nguyen et al., 2014). Recently, an agonistic C-C chemokine ligand of GPR15, named GPR15L, has been characterized (Ocón et al., 2017; Suply et al., 2017). GPR15L is expressed in colon and cervical epithelia and might play a role in mucosal immunity.

To discover novel endogenous GPR15 ligands, we screened a hemofiltrate-derived peptide library containing essentially all peptides and small proteins circulating in human blood in their final processed and physiologically relevant forms (Münch et al., 2014) for inhibitors of GPR15-mediated SIV infection. Multiple rounds of peptide separation and antiviral screening identified a C-terminal fragment of Cystatin C (named CysC95-146) as a potent and specific inhibitor of GPR15-dependent HIV and SIV infection. Cystatin C is a small (13 kDa) basic protein that is produced by all nucleated cells (Onopiuk et al., 2015) and represents the most abundant and potent extracellular inhibitor of cysteine proteases (Turk et al., 2012). It is found in virtually all tissues and body fluids and commonly used as a marker of renal function (Villa et al., 2005). Accumulating evidence suggests a role of Cystatin C in inflammation, neutrophil chemotaxis and resistance to bacterial as well as viral infections (Magister and Kos, 2013; Sokol and Schiemann, 2004; Xu et al., 2015; Zi and Yu 2018). We show that Cystatin C fragments preventing GPR15-dependent HIV and SIV infection are generated by proteases activated during antiviral immune responses. Unexpectedly, the recently discovered chemokine ligand of GPR15, GPR15L (Ocón et al., 2017; Suply et al., 2017) had no significant effect on viral entry. In addition, CysC95-146 prevented SIV and HIV-2 infection without interfering with GPR15L-mediated signaling. Our data support that naturally occurring Cystatin C fragments are capable of blocking GPR15-mediated primate lentiviral infection without interfering with the physiological signaling function of this GPCR.

## RESULTS

### Identification of a novel GPR15-specific SIV inhibitor

To identify novel ligands of GPR15, peptide libraries were generated from up to 10,000 liters of hemofiltrate derived from individuals with chronic renal failure by cation exchange separation followed by reverse-phase (RP) chromatography (Münch et al., 2014; Schulz-Knappe et al., 1997). Initially, the hemofiltrate was applied to a large cation exchange column and eluted using eight buffers with increasing pH ranging from 2.5 to 9.0. Subsequently, the resulting pH-pools (P1 to P8) were separated into ~40–50 peptide-containing fractions (F1 to F50) by RP chromatography. The final library comprised about 360 peptide fractions representing essentially the entire blood peptidome in a highly concentrated, salt-free and bioactive form (reviewed in Bosso et al., 2018; Münch et al., 2014). Screening of this peptide library using the infectious SIVmac239 molecular clone that efficiently utilizes GPR15 (Pöhlmann et al., 1999) and human osteosarcoma (GHOST) cells stably expressing CD4 and GPR15 (Mörner et al., 1999) identified several neighboring fractions from pH pool 4 that inhibited virus infection. Fractions 19 and 20 were selected for further purification because they blocked GPR15-mediated SIVmac infection by ~95% (Figure 1A) without causing cytotoxic effects or affecting CCR5- and CXCR6-dependent viral entry (data not shown).

**Figure 1.**
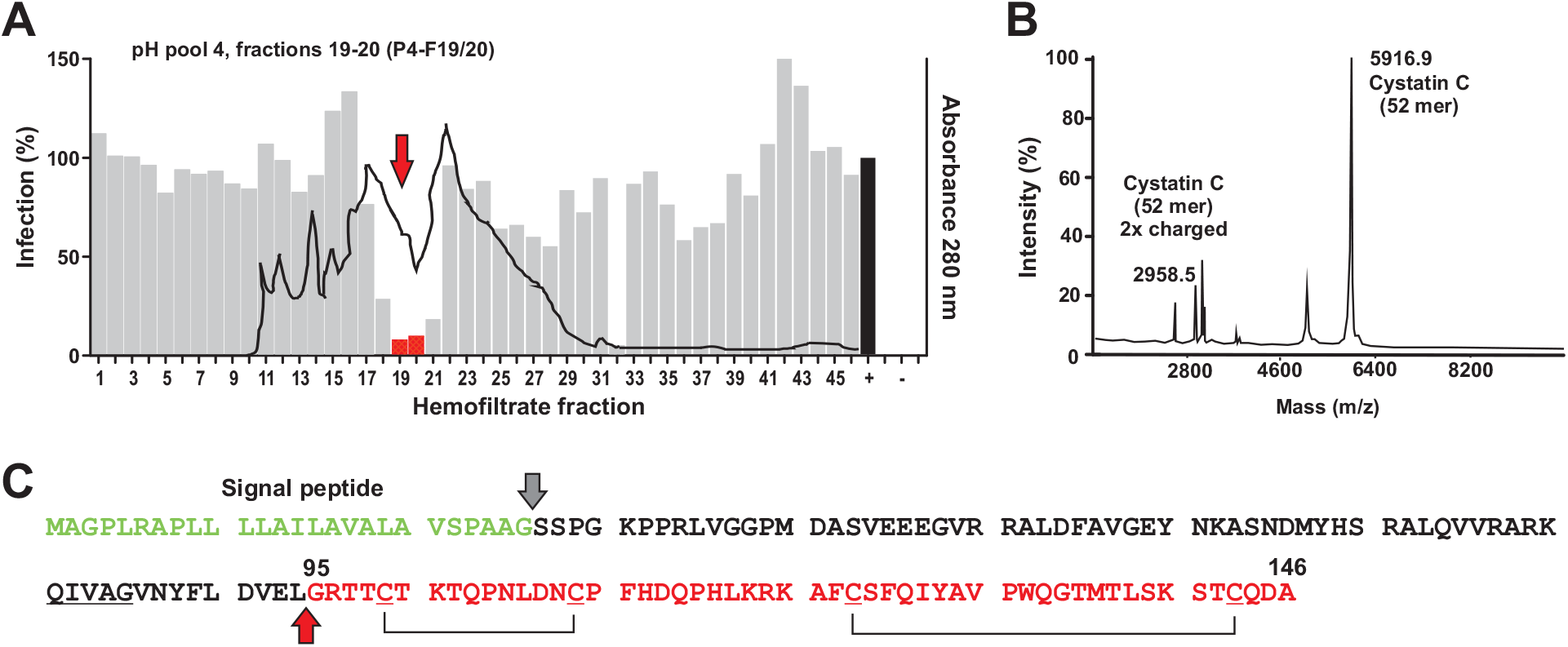
Identification of a C-terminal Cystatin C fragment inhibiting GPR15-mediated SIVmac infection. (**A**) The gray bars indicate the efficiency of SIVmac239 infection of GHOST-GPR15 cells in the presence of the hemofiltrate peptide library fractions compared to the absence of peptide (100%) and the black line indicates the peptide/protein elution profile. Fractions used for further peptide purification are indicated in red and highlighted by an arrow. + indicate infection in the absence of peptide, - shows uninfected cells. (**B**) MALDI-TOF spectrum of the active fraction obtained after the fifth round of purification. Sequence analyses identified the 52 C-terminal residues of Cystatin C. (**C**) Amino acid sequence of human Cystatin C. The signal peptide (green), the isolated peptide (red) and putative C-C bridges are indicated. The cleavage site to generate CysC95-146 is indicated by a red arrow.

MALDI-TOF mass spectrometry analysis of the inhibitory fraction obtained after five rounds of purification revealed a main m/z signal at 5.914 and some minor peaks (Figure 1B). The search in our (updated version) hemofiltrate peptide database (Richter et al., 1999) revealed that this signal corresponds to the 52 C-terminal amino acids (aa) of Cystatin C (aa 95 to 146) with an oxidation in Met136 (Figure 1C). Another m/z signal at 2.959 corresponded to the double-charged species of the same peptide. Cystatin C is expressed as a 146 aa precursor with a 26 aa signal peptide (GenBank accession number: AAA52164.1). It represents the most abundant and potent inhibitor of cysteine proteases in the human body and is an early marker of renal failure (Randers et al., 1998; Turk et al., 2012).

### A natural CysC fragment inhibits GPR15-dependent SIV and HIV infection

To verify that the identified peptide is responsible for the antiviral activity, we chemically synthesized the 52 C-terminal aa residues of Cystatin C (indicated in Figure 1C). The synthetic peptide, referred to as CysC95-146, inhibited GPR15-mediated SIVmac239 infection in a dose-dependent manner with a mean 50% inhibitory concentration (IC_50_) of ~0.5 μM (Figure 2A). Potent inhibition was confirmed for three divergent infectious molecular clones (IMCs) of SIVsmm (Figures 2B, S1), which naturally infects sooty mangabeys (Chahroudi et al., 2012) and frequently utilizes GPR15 as entry cofactor (Riddick et al., 2010). CysC95-146 had little if any effect on CCR5-or CXCR6-mediated virus infection, suggesting that it is specific for GPR15 (Figure 2B) and did not display significant cytotoxic effects (Figure 2C).

**Figure 2.**
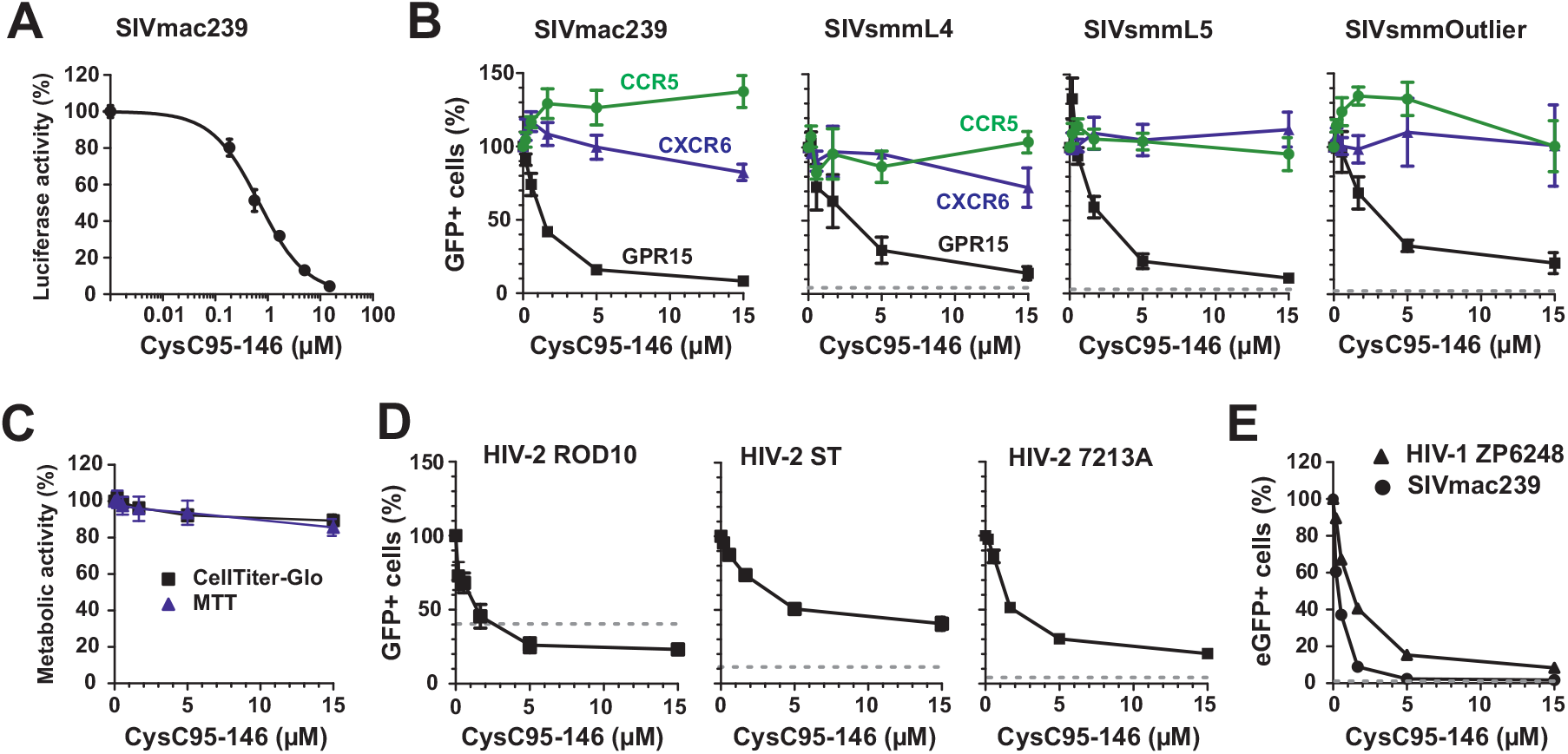
CysC95-146 specifically inhibits GPR15-mediated SIV and HIV infection. (**A**) GHOST-GPR15 cells were infected with a SIVmac239 luciferase reporter construct in the presence of CysC95-146. Experiments shown in all panels were performed at least in triplicate and curves show mean values (±SEM). (**B**) GHOST cells engineered to express GPR15, CXCR6 or CCR5 were infected with different SIV strains. Values show the percentage of virally infected (GFP+) cells in the presence of increasing concentrations of CysC95-146 compared to the percentage of infected cells obtained in the absence of peptide (100%). The dotted line indicates the percentage of eGFP+ cells obtained after infection of the parental GHOST cell line in the absence of peptide. (**C**) CysC95-146 is not cytotoxic. GHOST-GPR15 seeded in 96-well F-bottom plates were incubated with increasing amounts of peptide for 3 days at 37°C. Metabolic activity was analyzed by MTT and CellTiter-Glo assay. (**D, E**) Inhibition of (D) the indicated HIV-2 molecular clones or (E) HIV-1 ZP6248 by CysC95-146. In panel E, SIVmac239 is shown as positive control for comparison. Experiments were performed as described in (B). See also Figure S1.

SIVsmm is the precursor of HIV-2 (Sharp and Hahn, 2011), which is endemic in West Africa and has infected about one to two million people (Visseaux et al., 2016). To evaluate a possible role of CysC95-146 in HIV-2 infection, we examined its effect on infection by three HIV-2 strains: ROD10, representing a derivative of the first reported infectious HIV-2 clone (Clavel et al., 1986; Döring et al., 2016); ST, an attenuated HIV-2 strain with low *in vitro* cytopathic activity (Kong et al., 1988; Kumar et al., 1990) and 7312A, originally isolated from an individual from Côte d’Ivoire with dual HIV-1 and HIV-2 infection (Gao et al., 1994). HIV-2 ROD10 and ST belong to the group A of HIV-2 that is most widespread in the human population, while 7312A represents a recombinant form of groups A and B (Ibe et al., 2010). CysC95-146 inhibited all HIV-2 strains in a dose-dependent manner with IC_50_ values ranging from 1.3 to 7.0 μM (Figure 2D). However, maximal inhibition of HIV-2 entry did not exceed ~80%. One reason for this might be GPR15-independent infection of GHOST cells. Indeed, HIV-2 ROD10 infected the parental GHOST cell line that expresses low levels of CXCR4 but not GPR15 or other known entry cofactors with significant efficiency (Figures 2D; S1).

HIV-1 is less promiscuous in coreceptor usage than HIV-2. It usually utilizes only CCR5 during chronic infection and frequently evolves the ability to use CXCR4 during or after AIDS progression (Connor et al., 1997; Xiao et al., 1998). Some HIV-1 isolates, however, may also utilize GPR15 for productive infection and replication especially at high expression levels (Deng et al., 1997; Krumbiegel et al., 1999; Xiao et al., 1998). Interestingly, it has been reported that the transmitted/founder HIV-1 IMC ZP6248 is severely impaired in CCR5 and CXCR4 coreceptor usage but capable of infecting cell lines expressing alternative coreceptors including GPR15 (Jiang et al., 2011). CysC95-146 inhibited infection by the HIV-1 ZP6248 molecular clone in GPR15-GHOST cells with an IC_50_ of 1.1 μM (Figure 2E). Altogether, our results showed that CysC95-146 specifically inhibits GPR15 dependent SIV and HIV infection.

### The inhibitory CysC95-146 fragment binds to GPR15 expressing cells

To examine whether CysC95-146 specifically targets GPR15-expressing cells, we generated N- and C-terminal fusions of this GPCR with the enhanced green fluorescent protein (eGFP). N-terminal tagging with eGFP resulted in mislocalization of GPR15 and lack of cell surface expression (Figure S2A). In comparison, the C-terminally tagged GPR15-eGFP was efficiently expressed and showed proper localization at the cell surface (Figure S2B). For colocalization studies, we transfected HeLa cells with empty control or GPR15-eGFP vector and exposed the cells to Atto647-labeled CysC95-146. The antiviral peptide was readily detectable at the surface of GPR15-expressing HeLa cells but absent from control cells (Figure 3A). Quantitative analysis confirmed that CysC95-146 accumulates specifically on GPR15-eGFP-expressing cells (Figure 3B). Consistent with its proper subcellular localization, GPR15-eGFP-mediated CD4-dependent entry of SIV with an efficiency similar to the parental GPR15 protein (Figure 3C). These results are further evidence that CysC95-146 specifically targets GPR15 at the cell surface.

**Figure 3.**
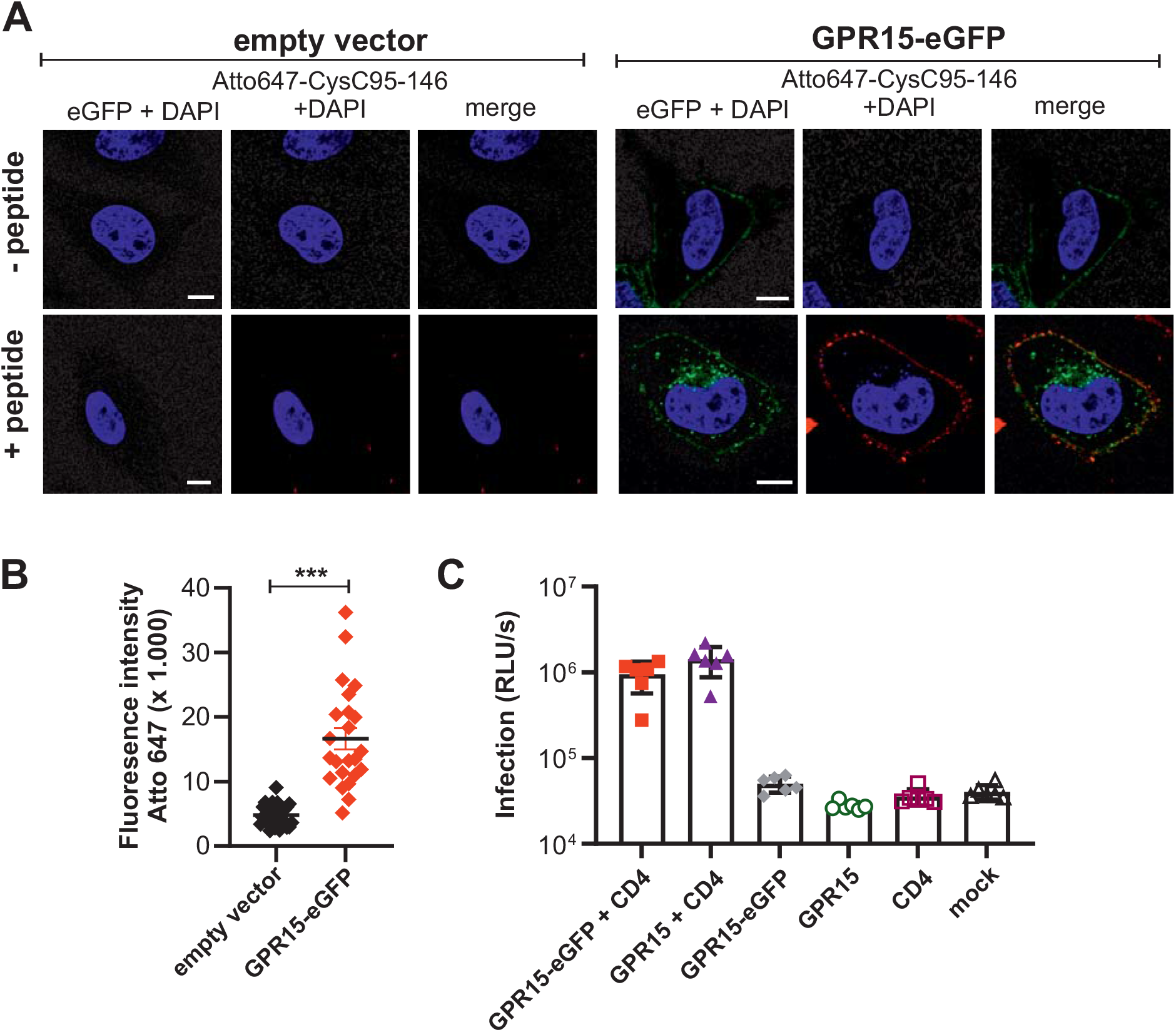
CysC95-146 interacts specifically with GPR15-expressing cells. (**A**) Hela cells were transfected with an eGFP-tagged GPR15 expression plasmid or a vector control and cultured for 2 days. Subsequently, cells were treated with Atto647-labeled CysC95-146. Excessive peptide was removed in several washing steps before staining the nuclei with Hoechst33342 (Merck) according to the manufacturer’s instructions. Samples were analysed by confocal microscopy using a LSM710 (Carl Zeiss). Scale bar indicates 10 μm. (**B**) Quantification of the Atto647 mean fluorescence intensity at the cell membrane from Atto-647-CysC95-146-treated samples shown in panel A. n=23-27 cells +/- SEM. ***, p<0.001 (Mann-Whitney test, unpaired t-test, non-parametric). (**C**) HEK293T cells transiently transfected with plasmids expressing the indicated receptors and/or an empty control vector and subsequently infected with a SIVmac239 Nano-Luciferase reporter virus. Infection rates were quantified as relative light units (RLU) per second. Results are displayed as means +/- SEM, n = 6. One representative of two biological repeats is shown; **, p<0.01 (Mann-Whitney test, unpaired t-test, nonparametric) See also Figure S2.

### A variety of C-terminal CysC fragments prevent GPR15-mediated lentiviral infection

To further examine the specificity of the antiviral activity of CysC95-146, we determined whether full-length Cystatin C and other C-terminal fragments also affect primate lentiviral infection. Our results showed that full-length Cystatin C displayed little inhibitory effect on SIVmac239 entry (Figure 4A). In contrast, N-terminal truncations of CysC95-146 by up to 12 amino acid residues (CysC107-146) (Figure 4A), as well as expansion by up to six residues (89-146) (Figure 4B) did not disrupt its antiviral activity. However, a peptide containing an additional ten residues at its N-terminus (CysC85-146) did not inhibit SIVmac entry into GPR15-GHOST cells (Figure 4B). Comprehensive analyses of a variety of C-terminal Cystatin C fragments revealed that residues 107 to 140 are sufficient for antiviral activity (Figure 4C). Further truncations at the N-or C-termini resulted in reduced activity or fully disrupted inhibition of GPR15-mediated SIVmac239 infection. Notably, some of the Cystatin C fragments analyzed, such as CysC107-146, were even more potent than CysC95-146 in inhibiting GPR15-mediated SIVmac infection (Figure 4C). Altogether, the results showed that a large variety of C-terminal Cystatin C fragments prevent GPR15-mediated SIVmac infection.

**Figure 4.**
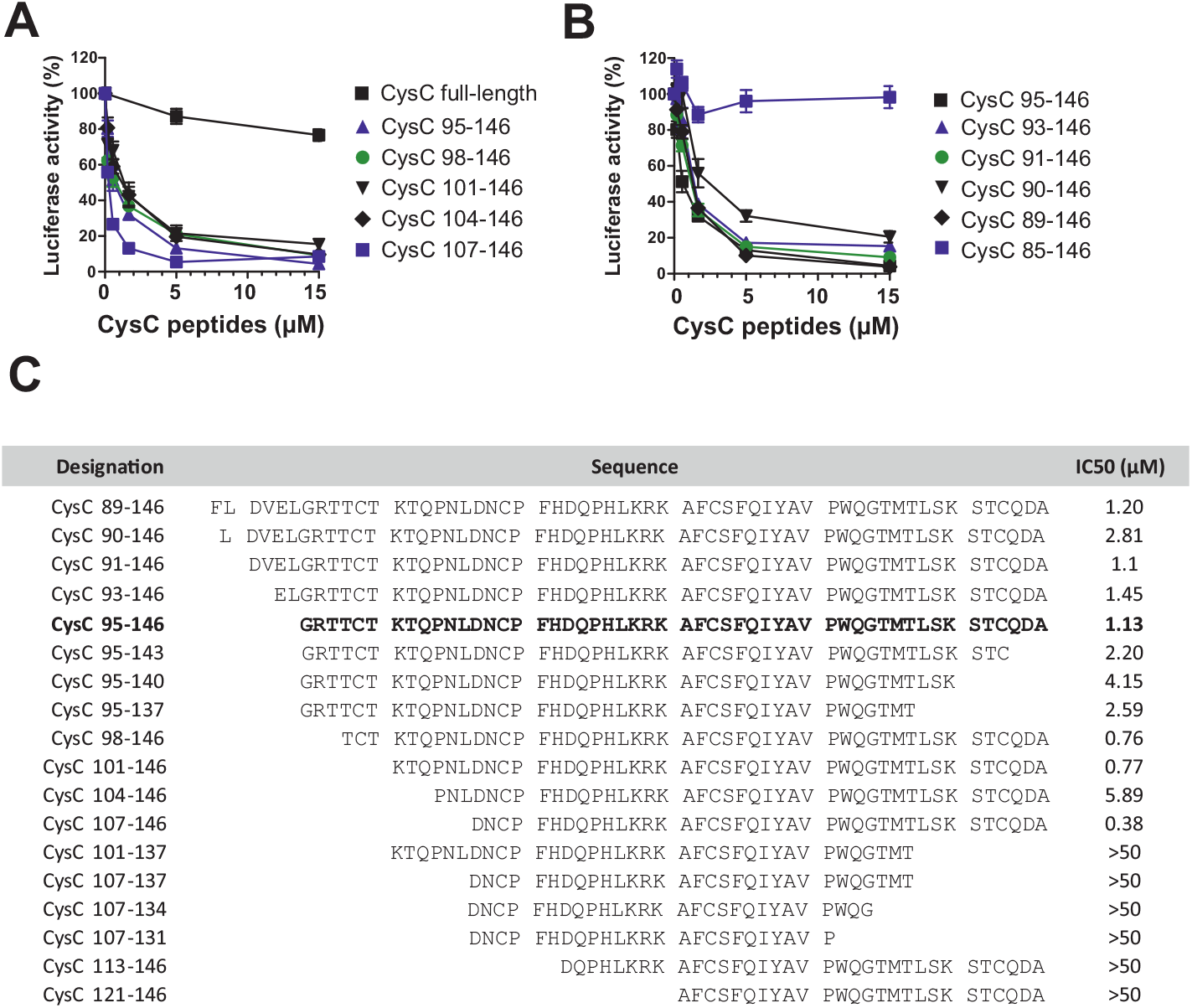
Anti-SIV activity of different C-terminal Cystatin C fragments. (**A, B**) Inhibition of SIVmac239 by (A) full-length Cystatin C and N-terminally truncated or (B) extended CysC peptides. Numbers refer to amino acid positions in the full-length Cystatin C sequence. GHOST GPR15 cells were treated with the indicated concentrations of the Cystatin peptides for 2h at 37°C before infection with a SIVmac239 Firefly-Luciferase reporter virus. Three days post infection, firefly-luciferase activities were analyzed. Experiments shown in all panels were performed at least in triplicates and curves show mean values (±SEM). (**C**) Overview of the amino acid sequences and antiviral activity of chemically synthesized CysC peptides analyzed. IC_50_ values were determined by infection of GHOST-GPR15 cells with a SIVmac239 luciferase reporter construct in the presence of increasing quantities of the indicated peptides.

### Proteolytic generation of antiviral CysC fragments

CysC95-146 was isolated from a hemofiltrate-derived peptide library, suggesting that it naturally circulates in the blood stream. To further examine this, we developed an MS-based method for the quantification of CysC95-146 in hemofiltrate (Figure S3A). These analyses showed that CysC95-146 was present at concentrations of ~10.7 ng/ml (236 pmol) in the original hemofiltrate fraction (Figures S3B, S3C). This concentration is lower than the IC_50_. However, hemofiltrate is significantly diluted compared to blood. In addition, material may have been lost during sample preparation and other C-terminal fragments are also antivirally active (Figure 4). In addition, Cystatin C levels are elevated in HIV-infected patients compared to healthy individuals (Neuhaus et al., 2010; Odden et al., 2007). Thus, our measurements most likely underestimate the concentrations of antiviral C-terminal Cystatin C fragments that can be achieved in vivo.

To examine the generation of antiviral peptides from Cystatin C, we treated the full-length protein with cathepsins C, D and G, as well as trypsin, chymase and Napsin A (Figure 5A). These proteases represent major components of the endo- and lysosomal protein degradation machinery (Turk et al., 2012; Zaidi and Kalbacher, 2008) and some of them are efficiently released from immune cells during infectious and inflammatory processes (Appelqvist et al., 2013; Yamamoto et al., 2012). Treatment of Cystatin C with cathepsin D, trypsin, pepsin, chymase and napsin A resulted in the generation of peptides with sizes similar to CysC95-146 (Figure 5A). Products obtained after digestion with cathepsin D, chymase and napsin A significantly inhibited GPR15-mediated SIVmac infection (Figure 5B). In addition, mass spectrometry of the digestion products confirmed the presence of several antivirally active Cystatin C fragments in cathepsin D, chymase and napsin A digested samples (Fig. 5C). Cathepsin D is an important component of the lysosomal protein degradation pathway in virtually all cells (Sun et al., 2013) and released from immune cells during inflammatory processes (Appelqvist et al., 2013). Chymase is a serine protease that is mainly produced by activated mast cells and elevated in some viral infections (Tissera et al., 2017). Napsin A is an aspartic proteinase that is abundantly expressed in normal lung and kidney tissue and a marker for some neoplasia (Bishop et al., 2010). Altogether, these results show that Cystatin C fragments which inhibit GPR15-mediated HIV and SIV infection are detectable in blood-derived human hemofiltrate and can be generated by proteases that are present and activated at sites of infection and inflammation.

**Figure 5.**
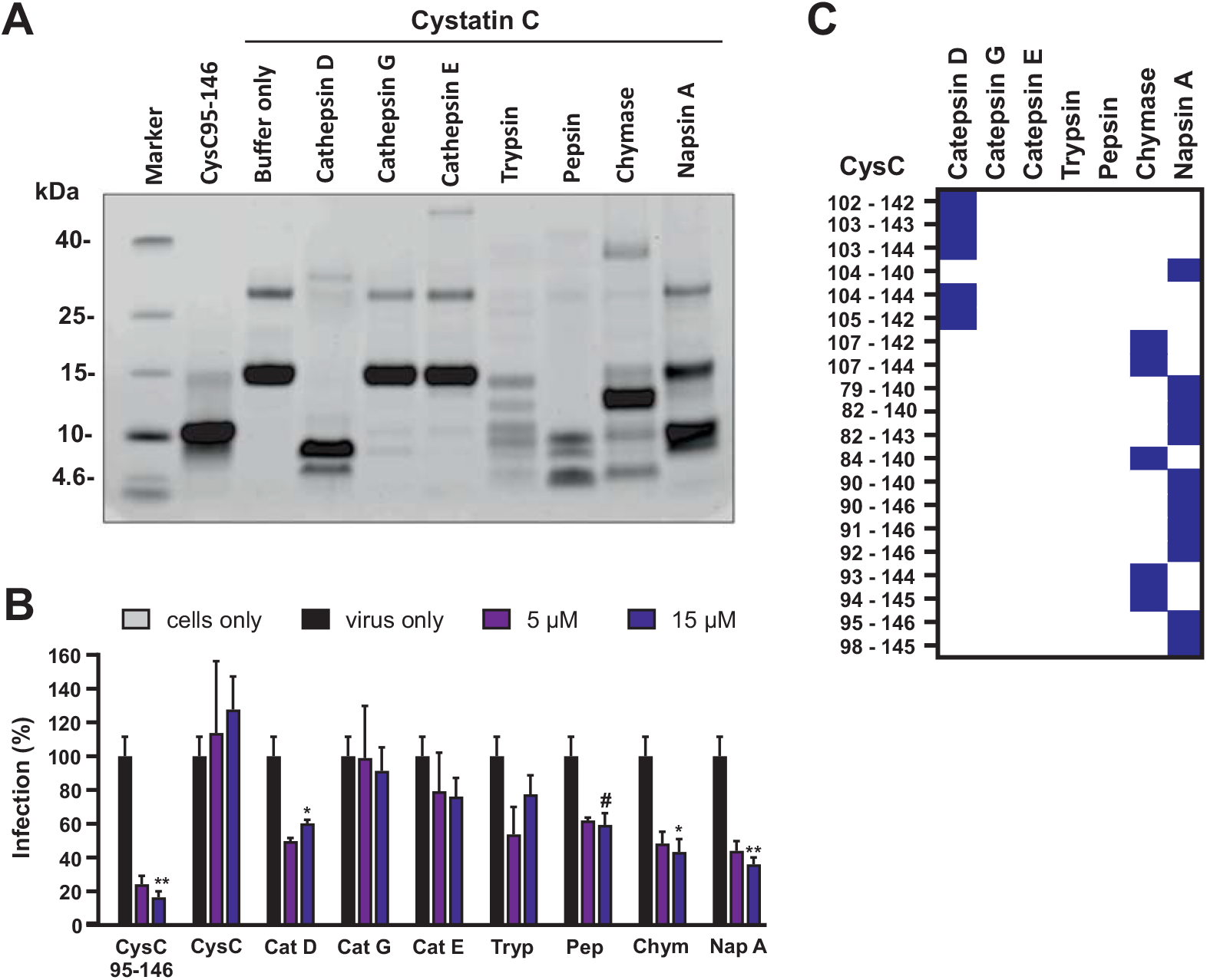
Treatment with various proteases generates antiviral Cystatin C fragments. (**A**) Human Cystatin C protein was digested with the indicated proteases. Digestion products were separated by SDS-PAGE and visualized by Coomassie Brilliant Blue staining. As controls, non-digested Cystatin C as well as the synthesized CysC95-146 were included. (**B**) Proteases and non-digested protein were removed by ultrafiltration with different kDa cut-offs for purification. GHOST-GPR15 cells were then treated with the digestion products or CysC95-146 or full-length Cystatin C and subsequently infected with a SIVmac239 luciferase reporter virus. Infection was measured at 3 dpi as described before. Results are displayed as means +/- SD of one experiment in triplicates. **, p<0.01 (Mann-Whitney test, unpaired t-test, nonparametric). (**C**) Heat Map visualization of identified Cystatin C fragments in samples digested with the indicated proteases by Mass spectrometry. See also Figure S3.

### GPR15L does not prevent SIV entry

For a long time, GPR15 had remained an orphan receptor but recently an agonistic chemokine ligand (named GPR15L) that modulates lymphocyte recruitment to epithelia has been identified (Ocón et al., 2017; Suply et al., 2017). It is well established that the chemokine ligands of the main entry cofactors of HIV-1, CCL5/RANTES and CXCL-12/SDF-1, inhibit CCR5-or CXCR4-mediated HIV-1 infection, respectively (reviewed in Verani and Lusso, 2002). Unexpectedly, GPR15L did not display an inhibitory effect on GPR15-mediated SIVmac infection (Figure 6A), although it induced downregulation of GPR15 from the surface of GHOST-GPR15 and CEM-M7 cells (Figures 6B; S4A). Receptor internalization was strongly reduced when the cells were kept on ice indicating that it mainly resulted from endocytosis and not from competition with the antibody used for staining. In contrast to GPR15L, CysC95-146 did not affect cell surface expression of GPR15 (Figures 6C; S4A). In addition, CysC95-146 did not induce calcium release in CHO cells engineered to express human GPR15 together with the promiscuous G-protein Gα16 (Suply et al., 2017), while GPR15 signaling was confirmed for GPR15L (Figure 6D). Altogether, the results showed that CysC95-146 prevents SIV infection without altering GPR15 cell surface expression, while downmodulation of GPR15 by GPR15L was insufficient to cause significant inhibitory effects on SIV entry.

**Figure 6.**
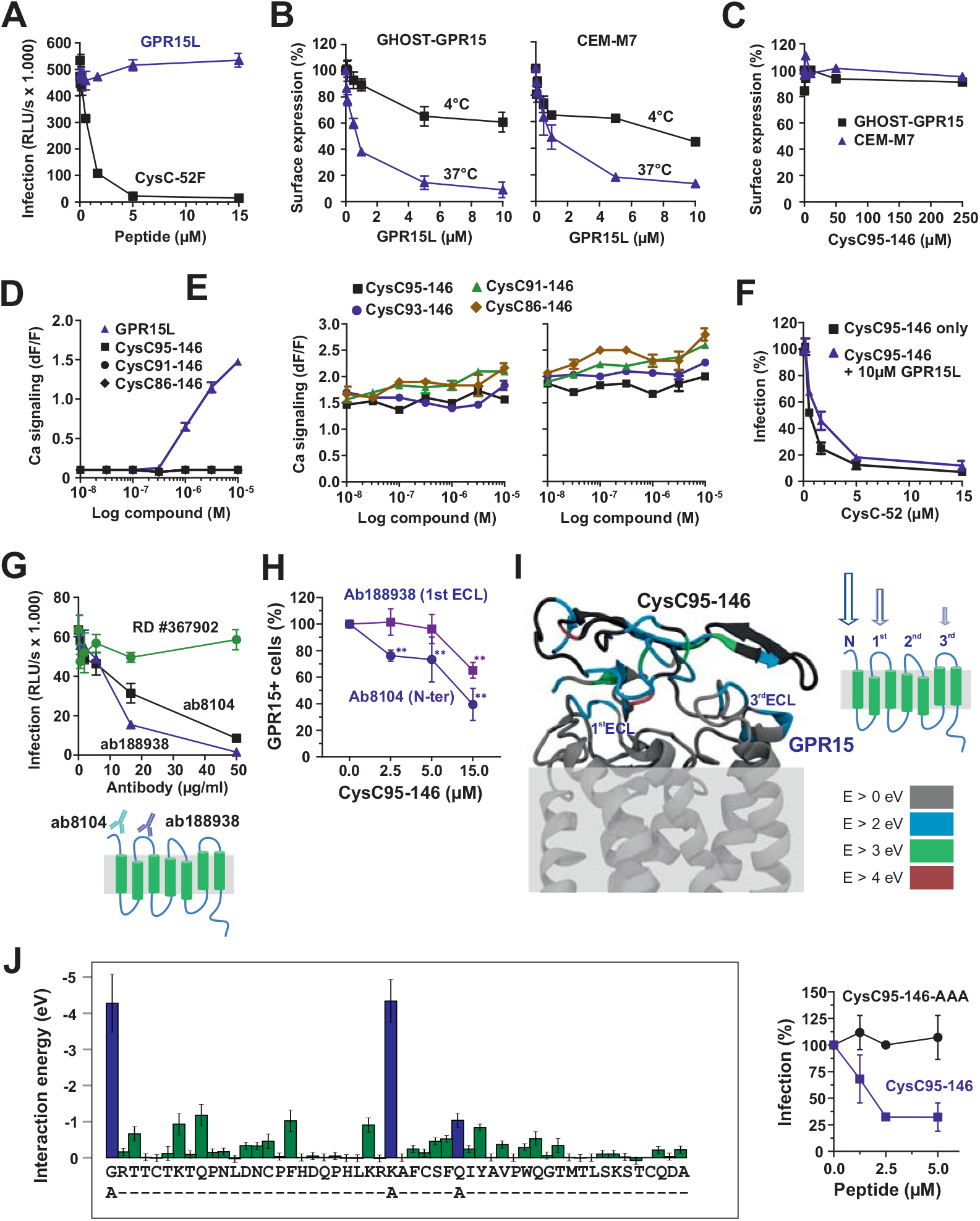
Effect of CysC95-146 and GPR15-L on GPR15 function. (**A**) GPR15L does not inhibit SIVmac239 infection. GHOST-GPR15 cells were preincubated with increasing amounts of GPR15L (Novartis) or CysC95-146 and subsequently infected with the SIVmac239 luciferase reporter virus. Three days post infection, infection rates were analyzed using a firefly luciferase reporter assay. Experiments shown in all panels were performed at least in triplicate and curves show mean values (±SEM). (**B, C**) GPR15-L (B) but not CysC95-146 (C) downmodulate GPR15 from the surface of GHOST-GPR15 and CEM-M7 cells. GHOST-GPR15 or CEM-M7 cells were preincubated with increasing amounts of GPR15L or CysC95-146 at 4°C (to prevent receptor internalization prior to staining with anti-GPR15 or isotype control antibody). GPR15 expression was analyzed by flow cytometry. (**D**) Cystatin C fragments do not induce GPR15-mediated calcium signaling. The effect of the indicated CysC fragments and GPR15L on calcium signaling was detected by measuring aequorin fluorescence in GPR15-expressing CHO-K1 cells. Each data point represents the mean relative light units ±SD over background fluorescence of quadruplicate measurements. (**E**) Cystatin C fragments do not interfere with the signaling function of GPR15L. The effect of the indicated concentrations of CysC fragments on GPR15L signaling was analyzed as described in panel D. (**F**) GPR15L does not enhance the antiviral activity of CysC95-146. GHOST-GPR15 cells were treated with increasing concentrations of CysC95-146 in the presence or absence of GPR15L. Cells were then infected with a SIVmac239 luciferase reporter virus. Inhibition of SIVmac infection was examined as described in panel A. (**G**) Anti-GPR15 antibodies ab8104 targeting the N-terminus and ab188938 targeting the 1^st^ extracellular loop of GPR15 but not RD #367902 (unknown binding site) inhibit SIVmac infection. GHOST-GPR15 cells were preincubated with increasing amounts of the indicated anti-GPR15 antibodies prior to infection with a SIVmac239 Luciferase reporter virus and analyzed as described in panel A. (**H**) GHOST-GPR15 cells were incubated with peptide to achieve the indicated concentrations of CysC 95-146 and either ab8104 targeting the N-terminus of GPR15 or ab188938 targeting the 1^st^ ECL of GPR15. Then, samples were washed and stained with a PE-conjugated secondary antibody and analysed by flow cytometry. Indicated frequencies of GPR15 positive cells were obtained by subtraction of background (secondary antibody only), followed by normalization to the no peptide samples. Shown are the mean values for n = 3 experiments ± SEM, * p < 0.05 (Welch’s t-test, unpaired). (**I**) Theoretically proposed binding of CysC95-146 with GPR15 based on reactive MD simulations (ReaxFF). In the model, the amino acid-resolved interaction energies are highlighted using the color coding shown in the lower right-hand scale. (**J**) Effect of amino acid changes of G69A, K94A and Q100A on the antiviral activity of CysC95-146. The left panel shows the predicted amino acid-resolved interaction energies and the position of amino acid changes whilst the right panel shows the effect of the parental and mutant CysC95-146 peptides on SIVmac239 infection in GHOST-GPR15 cells. Data shows mean values +/- SEM, ** p < 0.01, * p < 0.05 (Multiple t-test, unpaired, corrected for multiple comparisons using the Holm-Sidak method). See also Figure S4.

### CysC95-146 does not interfere with GPR15L signaling

The results outlined above suggested that CysC95-146 might inhibit GPR15-mediated viral entry without interfering with the physiological signaling activity of this GPCR. Indeed, calcium flux assays performed in the presence of constant quantities of GPR15L and increasing doses of various CysC fragments revealed that C-terminal CysC fragments did not reduce the signaling activity of GPR15L (Figure 6E). Conversely, GPR15L did not enhance the antiviral activity of CysC95-146 (Figure 6F). To obtain insights into the region(s) in GPR15 targeted by CysC95-146, we examined the ability of this antiviral peptide to compete with GPR15 antibodies. CysC95-146 competed with both Ab8104 targeting the extracellular N-terminus as well as (less efficiently) with Ab188938 interacting with the first extracellular loop (ECL1) (Figure 6G). These two antibodies also significantly inhibited SIVmac entry (Figure 6H). In contrast, Ab #367902 (R&D) that efficiently recognizes GPR15 in FACS-based assays and targets an unknown domain did not show significant effects on SIVmac infection (Figure 6G).

To further investigate the antiviral mechanism of CysC95-146, we performed molecular modelling on the interaction of CysC95-146 and GPR15 using structures obtained with reactive force field simulations. In agreement with the competition assays, the strongest interactions were predicted between CysC95-146 and the N-terminus of GPR15 (Figures 6I). To gain further insights into the interaction, the contribution of individual amino acids in GPR15 and CysC95-146 was analyzed in detail. We found that most amino acids in GPR15 (Figure S4B) and CysC95-146 (Figure 6J) stabilize the interaction by less than 2 eV. For GPR15, the strongest interactions were predicted for amino acid residues D2, E5, Y14 and T16, all located in the N-terminal part of this GPCR. In CysC95-146 only G95 and K120 stabilized the binding with >4 eV (Figure 6J) suggesting that these residues are important for interaction with GPR15. To test the theoretical predictions, we exchanged residues G95, K120 and Q126 in CysC95-146 to alanine. Indeed, these alterations fully disrupted the antiviral activity of CysC95-146 (Figure 6J). Altogether, our results suggest that the N-terminus of GPR15 is important for HIV-2 and SIV entry and the main target site for CysC95-146 interaction. In comparison, the second extracellular loop (ECL2) of GPCRs plays a key role in chemokine binding and signaling (Woolley and Conner, 2017). Thus, targeting of different surfaces on GPR15 might explain why the antiviral and agonistic functions of CysC95-146 and GPR15L did not interfere with one another.

### The antiviral activity of CysC95-146 is conserved in simian hosts of SIV

HIV-1 and HIV-2 are the result of at least thirteen independent zoonotic transmissions of SIVs from great apes or sooty mangabeys to humans that occurred in the last century (Sauter and Kirchhoff, 2019; Sharp and Hahn, 2011). In contrast, SIVs have infected non-human primate species for many thousands or even millions of years, possibly since primate speciation (Pandrea et al., 2008). GPR15 is a major SIV entry cofactor in the two best studied simian species naturally infected with SIV, i.e. sooty mangabeys and African green monkeys (Riddick et al., 2010, 2015). GPR15 also represents a major entry cofactor of SIVmac in macaques, the best established non-human primate model for AIDS in humans. To examine whether C-terminal Cystatin C fragments may play a role in SIV infection, we examined whether their ability to inhibit GPR15-mediated viral entry is conserved in monkeys. Cystatin C is the most ancestral member of the cystatin family of inhibitors of cysteine peptidases and found in many vertebrates (de Sousa-Pereira et al., 2014). Sequence alignments revealed that the C-terminus of human cystatin C is fully conserved in great apes and that the macaque ortholog differs only in two amino acids from its human counterpart (Figure 7A). We chemically synthesized the macaque CysC95-146 variant and found that it inhibits SIVmac239 infection as efficiently as the human version (Figure 7B). Thus, the antiviral activity of CysC95-146 is conserved in simian hosts of SIV infection.

**Figure 7.**
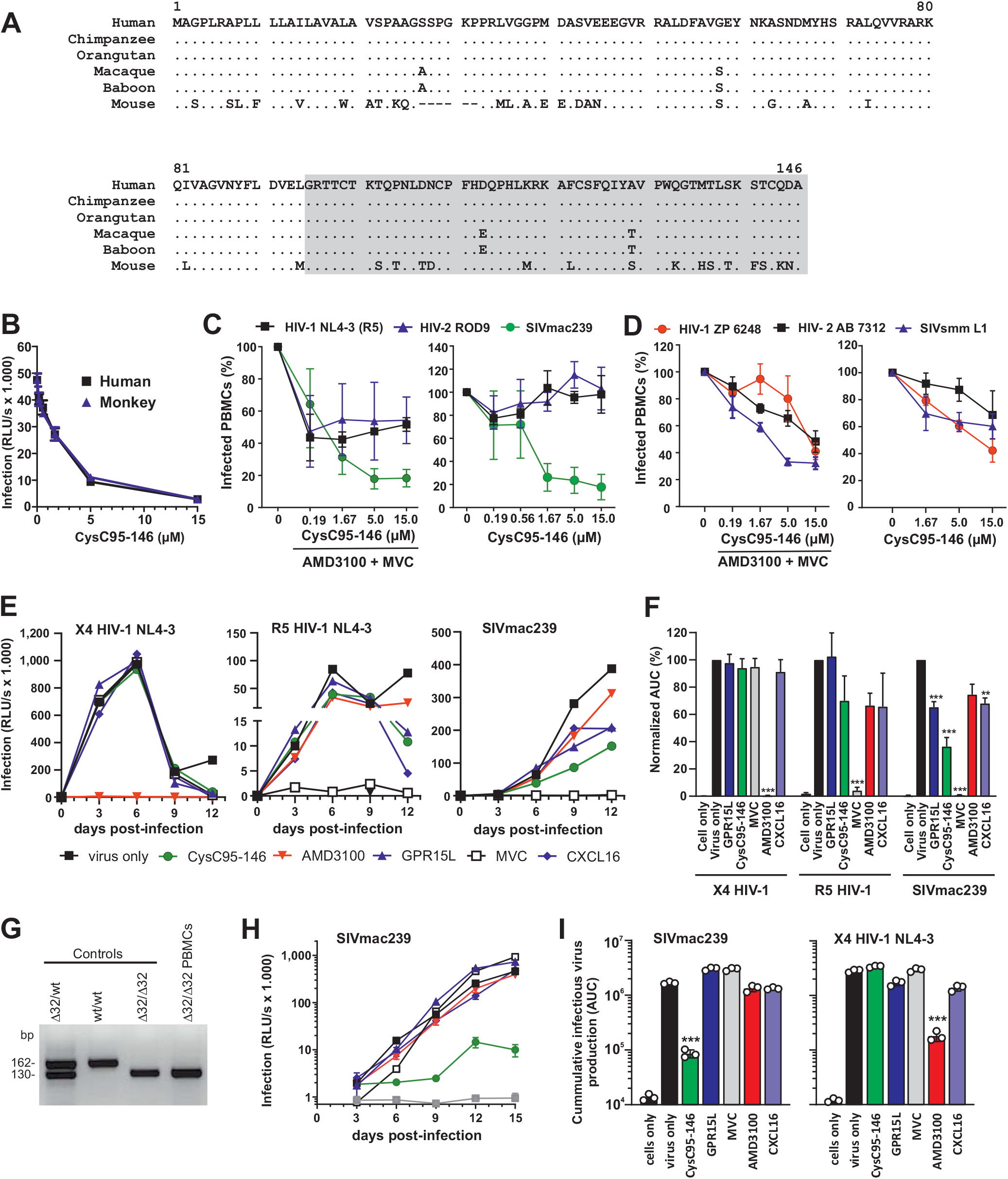
The antiviral activity of CysC95-146 is conserved in monkeys and affects HIV-2 and SIV infection in human T cells. (**A**) Alignment of Cystatin C amino acid sequences from the indicated species. Dots indicate identity to the human sequence and dashes gaps introduced to optimize the alignment. The region corresponding the CysC95-146 is shaded. (**B**) Antiviral activity of human and monkey-derived CysC95-146 peptides. GHOST-GPR15 cells were incubated with increasing amounts of human and monkey CysC95-146 for 2h at 37°C prior to infection with SIVmac239 F-Luc. At 3 dpi infection was analyzed via firefly luciferase reporter assay. The experiment was performed in triplicate. (**C, D**) Effect of CysC95-146 on SIV and HIV infection of primary human cells. Peripheral blood mononuclear cells (PBMCs) were isolated from buffy coats of three healthy donors, stimulated with IL-2/ PHA and treated with the indicated amounts of CysC95-146. Then, cells were infected with the indicated virus strains. 3 days post-infection, PBMCs were stained for p24 and analyzed by flow cytometry. Shown are the mean values for n = 3 donors +/- SEM and infection in cells obtained in the absence of peptide was set to 100%. *** p < 0.001, ** p < 0.01, * p < 0.05 (Multiple t-test, unpaired, corrected for multiple comparisons using the Holm-Sidak method). (**E**) CysC95-146 shows antiviral activity against GPR15-mediated SIVmac239 replication in human primary cells. To examine possible effects on spreading infection, we isolated and stimulated human PBMC as described before and treated those cells with the various compounds (CysC95-146, GPR15L, AMD3100, MVC, CXCL16) prior to virus exposure. Infectious virus production was determined by infection of TZM-bl indicator cells with PBMC culture supernatants obtained at different days post-infection. (**F**) Calculated Area-under-the-curve (AUC) for the virus replication data obtained in panel E. *** p < 0.001, ** p < 0.01, * p < 0.05 (Welch’s t-test, unpaired). (**G**) Verification of homozygous deletions in the *CCR5* gene of a Δ32/Δ32 PBMC donor. (**H**) Replication kinetics of SIVmac239 in Δ32/Δ32 PBMCs in the presence of various antiviral agents. Experimental details and symbols are provided in panel E. (**I**) Calculated area-under-the-curve (AUC) for the virus replication of SIVmac239 (see panel H) and X4 HIV-1 in Δ32/Δ32 PBMCs. See also Figure S5.

### Effect of CysC-F52 and GPR15L on SIV and HIV infection of primary T cells

To determine whether inhibition of GPR15-mediated HIV or SIV infection by CysC95-146 is relevant in primary viral target cells, we infected PHA-stimulated PBMCs from three human donors in the presence and absence of this peptide. Since PBMCs express various SIV and HIV coreceptors, we performed the experiment in the presence and absence of AMD3100 and Maraviroc, preventing CXCR4 and CCR5 mediated viral entry, respectively. FACS analyses allowed to determine the percentages of infected cells by intracellular p24 or p27 antigen staining (Figure S5A) and showed that PHA stimulation induced CCR5, GPR15 and CXCR6 cell surface expression (Figure S5B). CysC95-146 had no inhibitory effect on a CCR5-tropic derivative of HIV-1 NL4-3 and the CXCR4-tropic HIV-2 ROD9 molecular clone (Figure 7C). However, the peptide significantly reduced SIVmac239 infection of human PBMCs in both the presence or absence of additional inhibitors (Figure 7C). To further examine this, we analyzed the effects on three additional virus strains, i.e. the GPR15-tropic HIV-1 ZP6248 IMC, HIV-2 7312 and SIVsmm L1, which are capable of using CCR5, GPR15 and CXCR6 as entry cofactors (similarly to SIVmac239). CysC95-146 clearly reduced PBMC infection by all three HIV-1, HIV-2 and SIVsmm strains (Figure 7D) suggesting that GPR15-mediated entry contributes to primate lentiviral infection of primary CD4^+^ T cells.

To analyze possible effects on spreading infection, we pretreated PHA-stimulated human PBMC with the various inhibitors prior to virus exposure. Infectious virus production was determined by infection of TZM-bl indicator cells with PBMC culture supernatants obtained at different days post-infection. Predictably, AMD3100 blocked CXCR4-tropic HIV-1, while Maraviroc fully prevented CCR5-tropic HIV-1 replication (Figure 7E). In comparison, GPR15L, CysC-F52 and the CXCR6 chemokine ligand CXCL16 had no significant effect on CXCR4-or CCR5-tropic HIV-1 replication. Exposure of PBMCs to the GPR15-tropic HIV-1 ZP6248 strain did not result in significant replication precluding meaningful analysis of inhibitors (data not shown). However, GPR15L, CysC95-146 and CXCL16 all reduced replication of SIVmac239 in human PBMC cultures, albeit less efficiently than Maraviroc (Figure 7E). CysC95-146 was more effective than GPR15L and CXCL16 and suppressed infectious virus yield on average by ~60% (Figure 7F). FACS analyses revealed that CysC95-146 did not affect GPR15 cell surface expression while GPR15L reduced it by ~40% (Figures S5C, S5D). Unexpectedly, GPR15L induced significant downmodulation of CXCR4 (Figure S5C) and competed with the 12G antibody that targets ECL-2 of CXCR4 for binding to this GPCR (Figures S5E). However, in contrast to the small molecule inhibitor AMD3100, neither CysC95-146 nor GPR15L inhibited CXCR4-mediated HIV-1 infection (Figures S5F). Altogether, these results show that CysC95-146 suppresses GPR15-mediated primate lentiviral infection in primary human target cells and revealed an unexpected effect of GPR15L on CXCR4 cell surface expression.

We hypothesized that CysC95-146 may only reduce SIVmac239 replication by ~60% because this virus also utilizes CCR5 for entry into primary T cells. To examine this, we screened healthy uninfected individuals for the presence of the Δ32/Δ32 deletion in the *CCR5* gene that is present in about ~1% of Caucasians and disrupts functional CCR5 expression (Samson et al., 1996). We identified one donor containing homozygous deletions (Figure 7G) and performed infection experiments in Δ32/Δ32 PBMCs in the presence of various antiviral agents. Predictably, R5-tropic HIV-1 did not replicate in PBMCs lacking CCR5. Replication of SIVmac239 was almost entirely prevented by CysC95-146 but hardly affected by GPR15L, Maraviroc, AMD3100 or CXCL16 (Figure 7H). On average, CysC95-146 reduced production of infectious SIVmac239 by ~95% but had no inhibitory effect on X4-tropic HIV-1 (Figure 7I). *Vice versa*, AMD3100 reduced infectious yield of X4 HIV-1 by 94% but had no significant effect on SIVmac239. Thus, in the absence of CCR5, CysC95-146 prevents SIV replication in primary T cells almost entirely suggesting potent inhibition of GPR15 dependent virus entry.

## DISCUSSION

In the present study, we identified CysC95-146 and related C-terminal fragments of Cystatin C as effective and specific endogenous inhibitors of GPR15-dependent HIV and SIV infection. In contrast to CysC95-146, GPR15L, the recently discovered chemokine ligand of GPR15 (Ocón et al., 2017; Suply et al., 2017), displayed little if any inhibitory effect on HIV and SIV entry. This came as a surprise because the chemokine ligands of CCR5 and CXCR4 inhibit R5-or X4-tropic HIV-1 infection, respectively (Bleul et al., 1996; Oberlin et al., 1996; Walker et al., 1986). Notably, C-terminal fragments of Cystatin C prevent SIV and HIV-2 infection without interfering with GPR15L-mediated signaling activity of GPR15. In contrast, small molecule inhibitors of CCR5- and CXCR4-dependent HIV-1 entry, such as Maraviroc or AMD3100, antagonize chemokine signaling via these GPCRs (Donzella et al., 1998; Dorr et al., 2005). To our knowledge, CysC95-146 and related peptides are the first agents preventing GPCR-mediated infection by lentiviral pathogens without interfering with the signaling function of the corresponding chemokine receptor. Specific targeting of the detrimental function but not the signaling activity is of significant interest since GPCRs are involved in many physiological and pathological processes and the target of about 30% of all current drugs.

Cystatin C is produced by all nucleated cells, found in all tissues and body fluids, and represents the most abundant cysteine protease inhibitor (Villa et al., 2005). It is best known as a marker for renal failure. However, accumulating data also support an important role of Cystatin C in the immune response against various exogenous or endogenous pathogens (Magister and Kos, 2013; Neuhaus et al., 2010; Zi and Xu, 2018). The plasma levels in healthy individuals are about 0.1 μM. However, Cystatin C is induced in HIV infected individuals and reaches blood plasma levels up to 0.5 μM under conditions of renal failure, infection and inflammation (Bhasin et al., 2013; Longenecker et al., 2015; Randers et al., 1998). This concentration approximates the IC_50_ of antiviral Cystatin C fragments and it is conceivable that the local levels at sites of infection and inflammation might exceed the systemic plasma levels. In fact, we found that peptides blocking GPR15-mediated SIV entry are generated from Cystatin C by treatment with Cathepsin D, chymase and napsin A (Figure 5). These proteases are secreted by lysosomal exocytosis (Rodríguez et al., 1997) or via specialized secretory granules (Yamamoto et al., 2012) during immune responses and activated under acidic conditions. Acidification is a hallmark of inflammatory tissues (Okajima, 2013) and thought to play a key role in innate immunity (Rajamäki et al., 2013).

The generation of the active CysC fragments shows notable parallels to the generation of the CXCR4 antagonist and X4 HIV-1 inhibitor EPI-X4, which is generated from albumin by cathepsins D and E under acidic conditions (Zirafi et al., 2015). Albumin is more abundant than Cystatin C but the IC_50_ of EPI-X4 is also 10-fold higher than that of CysC95-146. Similarly, it has been reported that proteolytic processing of Chemokine (C-C motif) ligand 14 (CCL14), commonly also known as Hemofiltrate CC Chemokine 1 (HCC-1), by trypsin-like serine proteases generates a potent CCR5 agonist (CCL14[9-74]) that efficiently inhibits R5-tropic HIV-1 strains (Detheux et al., 2000; Münch et al., 2002). Similar to Cystatin C and albumin, the non-functional full-length CCL14 precursor is present at high concentrations in normal plasma. These results suggest that EPI-X4, CCL14[9-74] and active Cystatin C fragments are all preferentially generated at sites of infection and inflammation, where they might act locally to cooperatively inhibit CXCR4-, CCR5- and GPR15-mediated HIV or SIV infection, respectively. It is tempting to speculate that the combination of such endogenous inhibitors may have driven promiscuous coreceptor usage of SIVs that are most likely infecting primate species for millions of years (Compton et al., 2013; Gifford et al., 2008).

We have previously shown that structure-activity-relationship (SAR) studies allow to enhance the activity of endogenous peptides by several orders of magnitude and offer perspectives for therapeutic applications (Münch et al., 2007; Zirafi et al., 2015). For example, an optimized derivative of a natural 20-residue fragment of α(1)-antitrypsin that targets the gp41 fusion peptide of HIV-1 was safe and effective in human individuals (Forssmann et al., 2010). In addition, optimized derivatives of the endogenous CXCR4 antagonist Epi-X4 prevent atopic dermatitis and airway inflammation in preclinical mouse models (Harms et al., 2020). CysC95-146 is relatively large but tolerates truncations without loss of activity (Figure 4). We will perform structural and molecular modelling studies and SAR analyses to determine the minimal active size of C-terminal Cystatin C fragments and to increase their activity by rational design of derivatives predicted to interact more strongly with GPR15.

Our results show that CysC95-146 prevents GPR15-mediated HIV-2 and SIV infection, while GPR15L displayed little if any inhibitory activity although it induced downmodulation of GPR15 from the cell surface. It is known that both receptor removal from the cell surface as well as competitive inhibition by occupation of the interaction site(s) of the HIV envelope glycoprotein by chemokines might contribute to inhibition of CCR5-or CXCR4-dependent HIV-1 infection (Lobritz et al., 2013; Steen et al., 2009). Our data support that competition by C-terminal Cystatin C fragments is more effective than GPR15L-induced downregulation of GPR15 in inhibiting lentiviral infection in both GHOST indicator and primary CD4+ T cells suggesting that only a certain threshold is required for viral entry. Further structure-function analyses are required to fully elucidate the antiviral mechanism and the interaction(s) of CysC95-146 and GPR15L with GPR15. Our preliminary results from antibody competition assays and molecular modeling analyses suggest that CysC95-146 interacts most strongly with the N-terminal region of GPR15 (Figure 6I). The observed antiviral effect agrees with previous results showing that the N-terminal domains of CCR5 and CXCR4 are targeted by the HIV-1 envelope glycoproteins and play a key role in membrane fusion (Golding et al., 2005; Zhou et al., 2001). Consistent with published data on CCR5 and HIV-1, we found that antibodies targeting the N-terminus or ECL-1 of GPR15 prevented SIVmac239 infection (Figure 6G). GPR15L differs in GPCR interaction from prototype chemokines (Suply et al., 2017) and may interact with more C-terminal domains of GPR15. Differential GPR15 interaction sites also explain why CysC52 did not affect GPR15L-mediated signaling (Figure 6E) and *vice versa* the chemokine ligand did not enhance the inhibitory effect of the CysC95-146 fragment (Figure 6F).

Antiviral Cystatin C fragments may have some relevance in humans since GPR15 is a common coreceptor of HIV-2 that infects about one to two million people mainly in Sub-Saharan Africa and is also used by some HIV-1 strains. We show that CysC95-146 inhibits HIV-2 and the highly unusual HIV-1 ZP6248 strain not only in indicator cell lines but also in primary human cells (Figure 7). Our results also demonstrate that the antiviral activity of CysC95-146 is conserved in monkeys and support that the peptide inhibits SIV infection of primary human cells. HIV entered the human population only about a century ago and was hence clearly not a driving force in the evolution of endogenous peptides blocking GPR15-mediated entry. In comparison, SIVs infect non-human primate species for many thousand if not millions of years (Sharp and Hahn, 2011). GPR15 co-receptor usage is found in diverse groups of primate lentiviruses (Unutmaz et al., 1998) and most likely represents an ancient function. Thus, it is tempting to speculate that the evolution of inhibitors of GPR15-mediated viral entry by inflammation and infection-associated proteases might have been driven by ancient primate lentiviruses. Finally, our results suggest that GPR15 allows SIV to replicate in Δ32/Δ32 PBMCs cells in the absence of CCR5 (Figure 7G-I). In humans, the Δ32/Δ32 genotype is associated with a reduced risk of the acquisition of HIV-1 infection via the sexual route (Liu et al., 1996; Samson et al., 1996). Notably, sooty mangabeys, the original host of SIVsmm/HIV-2, also frequently lack functional CCR5 expression (Riddick et al., 2010). However, the prevalence of natural SIVsmm infection was not significantly reduced in animals lacking functional CCR5 most likely due to efficient coreceptor usage of GPR15 and CXCR6.

Our identification of CysC95-146 provides proof of concept that some ligands of GPCRs can block pathogens without interfering with their physiological signaling function. Notably, this is not the case for inhibitors of CCR5- and CXCR4-mediated HIV infection and this precludes e.g. usage of AMD3100 for the treatment of chronic diseases since proper CXCR4 signaling is critical for many physiological processes. After the CXCR4 antagonist EPI-X4 (Zirafi et al., 2015), CysC95-146 is another example of the proteolytic generation of a peptidic virus inhibitor from an abundant precursor protein by proteases that are activated under acidic conditions. It is conceivable that generation of anti-microbial effects by proteolytic generation of abundant precursors might provide a more effective and rapid means to generate innate immune effectors than *de novo* synthesis. Further studies to clarify whether generation of antimicrobial molecules by proteolysis of abundant precursor proteins by proteases activated during infection and inflammation represents a common concept of innate immune defense seem warranted.

## ACKNOWLEDGMENTS

A number of reagents were obtained through the NIH AIDS Reagent Program. NP, LS, AR, SW, MH, RG, CJ, TJ, JM and FK are supported by CRC 1279 of the Deutsche Forschungsgemeinschaft (DFG). KMJS is supported by a Mari-Sklodowska Curie fellowship (VIAR) and BHH is supported by R01 AI 114266 and UM1 AI 126620. The Alexander von Humboldt Foundation is gratefully acknowledged for financial support to A.R. (postdoctoral fellowship 3.2-KUB/1153731 STP).

## AUTHOR CONTRIBUTIONS

A.B. and M.H. performed most experiments. A.R. and L.S. purified CysC95-146 and N.P. synthesized peptides. A.R. and S.W. performed mass spectrometry and M.H. Ab competition experiments. C.S. generated proviral and GPCR expression constructs. Y.Y. and R.C. performed some infection experiments and B.H.H. provided viral constructs. C.J. and T.J. performed molecular modeling analyses. K.S. provided GPR15L and examined GPCR signaling. W.G.F. provided the HF-derived peptide library and J.M. reagents and expertise. F.K. supervised the study and wrote the initial draft of the manuscript. All authors have seen and approved the manuscript.

## STAR METHODS TEXT

### LEAD CONTACT AND MATERIALS AVAILABILITY

Further information and requests for resources and reagents should be directed to and will be fulfilled by the Lead Contact, Frank Kirchhoff. All unique/stable reagents generated in this study are available from the Lead Contact without restriction.

### EXPERIMENTAL MODEL AND SUBJECT DETAILS

#### Ethical statement for human samples

Experiments involving human blood and CD4+ T cells were reviewed and approved by the Institutional Review Board (i.e. the Ethics Committee of Ulm University). Individuals and/or their legal guardians provided written informed consent prior to donating blood. All human-derived samples were anonymized before use. The use of established cell lines did not require the approval of the Institutional Review Board.

#### Cell lines

GHOST BOB (GPR15), Bonzo (CXCR6), CCR5 and parental cells were obtained from the NIH AIDS Reagent Program. GHOST cells were maintained in in Dulbecco’s modified Eagle medium (DMEM) supplemented with FCS (10% (v/v)), L-glutamine (2 mM), streptomycin (100 mg/mL) and penicillin (100 U/mL), geneticin G418 (500 μg/ml), 100 μg/ml hygromycin and 1 μg/ml puromycin. The GHOST parental cell line is puromycin sensitive, so medium was not supplemented with Puromycin. Cells were cultured at 37°C, 90% humidity and 5% CO2. The cells were regularly split 1:10 or 1:20 twice a week. HEK293Tcells were provided and authenticated by the ATCC. TZM-bl cells were provided and authenticated by the NIH AIDS Reagent Program, Division of AIDS, NIAID. HEK293T, TZM-bl and Hela cells were maintained in Dulbecco’s modified Eagle medium (DMEM) supplemented with FCS (10%), L-glutamine (2 mM), streptomycin (100 mg/mL) and penicillin (100 U/mL). Cells were cultured at 37°C, 90% humidity and 5% CO2. CEM-M7 cells were cultured in RPMI 1640 medium with 10% fetal bovine serum, 1% penicillin-streptomycin, 1% 200 mM L-glutamine and 500 μg/ml G418.

#### Primary blood cells

Buffy coats (lymphocyte concentrates from 500 ml whole blood) were obtained from the blood bank (Ulm) and diluted 1:3 with PBS. Ficoll separating solution was overlaid with the diluted blood and centrifuged at 1,600 x g for 20 min without breaks. The white interface layer formed by peripheral blood mononuclear cells (PBMCs) was transferred into a fresh tube and washed twice with PBS. After separation and washing 1 x 106cells/ml were cultured in supplemented RPMI-1640 with 1 μg/ml PHA and 10 ng/ml IL-2 for 3 days.

### METHOD DETAILS

#### Generation and screening of HF libraries

Fractions of a hemofiltrate-derived peptide library generated as described (Zirafi et al., 2015) were tested for their ability to suppress SIVmac239 infection in GHOST-GPR15 cells. Cells were seeded in flat-bottomed 96-well dishes, cultured overnight, and incubated with the peptide for 2 h before infection with an SIVmac239 construct containing the firefly luciferase reporter gene in place of the *nef* gene. Forty-eight hours later, the cells were lysed with Cell Culture Lysis Reagent (catalog no. E153A; Promega) and relative light units (RLU) were determined using the luciferase assay system (Promega).

#### Purification of CysC95-146

The peptide was purified from a 1000 L HF-equivalent sample (308 mg) of the active fraction CEX-Pool 4/RP-Fr19-20 (peptide bank HF020802), by a combination of reversed-phase and cation-exchange HPLC techniques followed by biological testing of collected fractions after every purification step. The sequence of chromatographic steps was: A) Source RPC15 column of dimensions 20 x 250 mm; flow rate: 13 mL/min; gradient (min/%B): 0/2, 5/20, 20/30, 60/40, 70/60, 75/100, being A: 0.1% TFA and B: 0.05% TFA in 80% ACN; detection wavelength: 280 nm. B) Source RPC15 column of dimensions 20 x 250 mm; flow rate: 13 mL/min; gradient (min/%B): 0/2, 10/20, 58/32, 65/40, 68/60, and 75/100; detection wavelength: 280 nm. C) Source RPC15 column of dimensions 10 x 125 mm; flow rate:3 mL/min; gradient (min/%B): 0/5, 10/20, 60/28, 70/40, 68/60, and 80/100; detection wavelength: 280 nm. D) Grace Vydac 218TP54 RP18 column of dimensions 4.6 x 250 mm; flow rate: 1 mL/min; gradient (min/%B): 0/5, 5/26, 65/31, 70/36, and 75/100; detection wavelength: 225 nm. E) PolySULFOETHYL A cation-exchange column of dimensions 4 x 150 mm; flow rate: 0.7 mL/min; gradient (min/%B): 0/0, 30/30, 45/50, and 55/100, being A: 10 mM phosphate + 20% acetonitrile, pH 3.2, and B: 10 mM phosphate + 1M KCl + 20% acetonitrile pH 3.2; detection wavelength: 220 nm.

#### MALDI-TOF analysis of the active fraction

The molecular mass measurement of the biologically active fraction after the latest purification step (cation-exchange HPLC) was performed with a Voyager-DE Pro matrix-assisted laser desorption/ionization time-of-flight mass spectrometry (MALDI–TOF–MS) device (PerSeptive Biosystems, Framingham, MA, USA). The matrix solution was prepared with α-cyano-4-hydroxycinnamic acid dissolved at 5 mg/mL in mass buffer (0.1% TFA in 1:1 acetonitrile/water solution). The chromatographic fraction was desalted with an RPC18 cartridge (Waters, USA) prior to MS measurement. One microliter of the sample solution and matrix solution were mixed on a 100-well stainless steel MALDI plate. Measurements were performed in linear mode. Positive ions were accelerated at 20 kV, and up to 100 laser shots were automatically accumulated per sample position. Voyager RP BioSpectrometry Workstation version 3.07.1 (PerSeptive Biosystems, USA) was used as control and visualization software.

#### Synthetic peptides

Solid phase peptide synthesis was carried out using conventional Fmoc chemistry. Fmoc-protected amino acids with protected side chains were purchased from Novabiochem (Merck KGaA, Darmstadt, Germany). Synthesis was carried out using a preloaded Wang resin (Novabiochem) in a 0.1 mmol scale on a Liberty blue microwave peptide synthesizer (CEM Corporation, Matthews, NC, USA). After coupling reactions, deprotection and cleavage from the resin the raw peptides were precipitated and purified by reversed-phase HPLC. The correct molecular mass of the obtained purified synthetic peptides was analyzed by electrospray (ESI) and MALDI-mass spectrometry and their purity of 95% was confirmed by analytical HPLC. Human Cystatin C protein (purified, from human urine, BML-SE479-0100) was obtained from Enzo Life Sciences.

#### Quantitation of CysC95-146 in hemofiltrate by nanoLC-ESI-iTRAP-Orbitrap

CysC95-146 amount in hemofiltrate was estimated from the chromatographic fraction CEX-Pool 4/RP-Fr19-20 of the peptide bank HF020802, where the biological activity was initially found. For disulfide bridge reduction and carbamidomethylation, the standard (synthetic CysC95-146) and the HF sample were incubated with 50 mM NH_4_HCO_3_ + 5 mM DTT at RT for 20 min. Subsequently, iodoacetamide was added up to 50 mM to the samples and incubated at 37°C for 20 min. The reaction was stopped (quenching) with 10 mM DTT. The injection volume in the LC-MS/MS system was 15 μL. The amounts of injected peptide standards were (pmol) 62.5, 125, 250, 500 and 1000.

Samples were measured using an LTQ Orbitrap Velos Pro system (Thermo Fisher Scientific) online coupled to an U3000 RSLCnano (Thermo Fisher Scientific) as described (Mohr et al., 2015) with the following modifications: Separation was carried out using a binary solvent gradient consisting of solvent A (0.1% FA) and solvent B (86% ACN, 0.1% FA). The column was initially equilibrated in 5% B. In a first elution step, the percentage of B was raised from 5 to 15% in 5 min, followed by an increase from 15 to 40% B in 30 min. The column was washed with 95% B for 4 min and re-equilibrated with 5% B for 19 min. Ion chromatograms were visualized by XCalibur Qual Browser (Thermo Fisher Scientific, Bremen, Germany). For the identification of the CysC-F52 in the samples, the total ion chromatograms were filtered by searching the m/z range corresponding to the m/z value of the most intense signal (± 20 ppm) according to the theoretical isotopic distribution at z=6. The peak areas were calculated by using the default parameters of the software. The peak areas were exported and the linear regression calibration curve (peak area vs standard amount) was constructed in Microsoft Excel 2016.

#### Molecular clones of HIV and SIV

Generation of the SIVmac239 proviral construct carrying a functional *nef* gene followed by an IRES element and the eGFP cassette has been described previously (Schindler et al. 2006). A similar approach was used to generate the replication-competent SIVmac239 F-Luc construct. Therefore PCR amplification with flanking primers introducing MluI and XmaI restriction sites of the IRES-FLuc cassette from an HIV-1 IRES-FLuc reporter construct was used and therewith the IRES-eGFP cassette in SIVmac239 proviral construct replaced. The integrity of the PCR-derived insert was confirmed by sequencing. See Key reagent table for details on additional proviral HIV-1 and SIV constructs.

#### Virus stocks

Virus stocks were generated by transient transfection of HEK293T cells using the calcium-phosphate precipitation method or the TransIT®-LT1 Transfection Reagent (Mirus). One day before transfection, 0.65 x 106 HEK293T cells were seeded in 6-well plates (Greiner Bio-one, Frickenhausen, Germany). At a confluence of 50-75% cells were used for transfection. For the calcium-phosphate precipitation method, 5 μg DNA was mixed with 13 μl 2 M CaCl2 and the total volume was made up to 100 μl with water. This solution was added drop-wise to 100 μl of 2x HBS. The transfection cocktail was vortexed for 5 sec and added drop-wise to the cells. The transfected cells were incubated for 8-16 h before the medium was replaced by fresh supplemented DMEM. Transfection with TransIT®-LT1 Transfection Reagent was performed according to the manufacturer’s recommendation. 48 h post transfection, virus stocks were prepared by collecting the supernatant and centrifuging it at 1300 rpm for 3 min. SIVmac239 F-Luc virus stocks were additionally concentrated by ultracentrifugation. Virus stocks were stored at – 80°C.

#### Cell viability

GHOST Bob cells were seeded in 96-well F-bottom plates at a density of 10,000 cells per well one day prior to the experiment. Cells were washed once in PBS and incubated with increasing amounts of CysC95-146 for 3 days at 37°C. Metabolic activity was analyzed by MTT and CellTiter-Glo assay (Promega) according to the manufacturer’s recommendations.

#### Infectivity Assays

Effect of CysC95-146 and derivatives and GPR15L on GPR15-dependent SIVmac239 infection. GHOST Bob (GPR15) cells were seeded in 96-well plates at a density of 10,000 cells per well. The next day, cells were treated with Cystatin C derived peptides or GPR15L at the indicated concentrations and incubated for 1 – 2 hours at 37°C, 90% humidity and 5% CO2. Then, cells were infected with SIVmac239 F-Luc previously produced by transient transfection of HEK293T cells. Three days post infection, viral infectivity was determined using a Firefly-Luciferase Assay kit from Promega as recommended by the manufacturer. Firefly-Luciferase activities were quantified as relative light units (RLU) per second with an Orion Microplate luminometer (Berthold).

#### Co-receptor usage

GHOST-GPR15, CCR5 and CXCR6 cells were seeded 24h prior with a density of 6,000 cells/well in 96 well F-bottom plates. Cells were preincubated with increasing amounts of CysC95-146 for 2h at 37°C prior to virus addition. 3dpi, infection rates were analyzed by measuring GFP-positive cells via flow cytometry.

#### Inhibition by CysC95-146

GHOST-GPR15 cells were seeded 24h prior to infection at a density of 6,000 cells/well in 96-well F-bottom plates. Cells were incubated with increasing amounts of CysC95-146 from humans or agm/mac for 2h at 37°C prior to infection with SIVmac239 F-Luc. Three days post-infection, viral infectivity was determined using a Firefly-luciferase Assay kit from Promega as recommended by the manufacturer. Firefly-luciferase activities were quantified as relative light units (RLU) per second with an Orion Microplate luminometer (Berthold).

#### Infection of primary blood cells

1 million cells were incubated with various amounts of peptide or compound for 1 – 2 at 37°C. Then, cells were infected with virus stocks previously generated by transient transfection of HEK293T cells with the respective pro-viral constructs. PBMCs were cultured in the presence of 10 ng/ml IL-2 in RPMI-1640 supplemented with FCS (10% (v/v), L-glutamine (2 mM), streptomycin (100 mg/mL) and penicillin (100 U/mL). 3 days post infection cells were used for FACS analyses. Infected cells were first washed with 500 μl FACS buffer (PBS with 1% (v/v) FCS) and stained with 50 μl FACS buffer containing 1 μl FITC-conjugated p24 antibody. The cells were incubated for 30 min at 4°C and then washed with 1 ml FACS buffer to remove unbound antibody. Then, cells were fixed with 200 μl FACS buffer containing 2% PFA and incubated for 30 min at 4°C. PBMCs were gated based on forward and side scatter characteristics, followed by exclusion of doublets and then by the respective marker expression. Data were generated with BD FACS Diva 6.1.3 Software.

#### Replication kinetics in PBMCs

0.75 million cells were transferred into FACS tubes, washed twice in PBS and incubated with 10 μM GPR15L, 10 μM CyC95-146, 15 μM Maraviroc, 10 μg/ml AMD3100 or 500 ng/ mL CXCL16 in RPMI-1640 plus supplements for 1 hour at 37°C. Then, cells were infected with virus stocks previously generated by transient transfection of HEK293T cells with the respective pro-viral constructs. 16 hours post-infection, cells were washed and transferred to a 96 U-Well plate. At the indicated time points, cells were spun down and ~80% (v/v) of supernatants of the PBMC cultures were aspirated and frozen at −80°C. Medium was replaced with fresh RPMIxxx supplemented with 10 ng/mL IL-2. CysC95-146, GPR15L and the remaining compounds were added again to achieve the concentrations stated above.

#### Infectious virus yields from PBMCs

Infectivity of virions produced in primary cells in the presence and absence of CysC95-146, GPR15L, AMD3100, MVC, CXCL16. To determine the infectivity of virions produced in infected human and rhesus primary blood cells, TZM-bl cells were seeded in 96-well plates at a density of 10,000 cells/well and infected after overnight incubation with the supernatants collected from the PBMC cultures. Three days p.i., viral infectivity was determined using a galactosidase screen kit from Tropix as recommended by the manufacturer. β-Galactosidase activities were quantified as relative light units (RLU) per second with an Orion Microplate luminometer (Berthold).

#### GPR15 antibody competition assay

50,000 GHOST-GPR15 cells were counted, washed in FACS buffer (1x PBS with 1% FCS, 4°C) and transferred to FACS tubes. Cells were centrifuged at 4°C and 1,300 rpm (350 x g). Supernatants were discarded and cell pellets were resuspended in 100 μL master mix containing anti-GPR15 antibodies (ab8104 and ab188938) and CysC 95-146. Indicated concentrations of CysC 95-146 were achieved during this inoculation step. Antibody concentrations were 10 ng/mL. Cells were incubated at 4°C for 1 hour and washed 3 times in FACS buffer. Then, staining with a secondary antibody (Goat anti-rabbit IgG H&L PE, ab97070) was performed. For this, samples were incubated with secondary antibody at 10 ng/mL for 30 minutes at 4°C. After staining with the secondary antibody, cells were washed 3 times in FACS buffer, fixed in 4% PFA (1 h, 4°C) and kept at 4°C until analysis by flow cytometry. Cells were gated based on forward and side scatter characteristics, followed by exclusion of doublets and then by the respective marker expression. Data were generated with BD FACS Diva 6.1.3 Software before analysis with FlowJo 8.8 Software (Treestar).

#### Receptor downmodulation

GHOST-GPR15 or CEM-M7 cells were preincubated with increasing amounts of GPR15L or CysC95-146 for 30 min at either 37°C or 4°C (to prevent receptor internalization) prior to staining with anti-GPR15 or isotype control antibody. GPR15 expression was analyzed by flow cytometry. which Ab was used by Andrea in this experiment?

#### Flow cytometric analysis

Human PBMCs were isolated from whole blood as described before. 250,000 cells were incubated with 10 μM of CysC95-146 or GPR15L in RPMI-1640 supplemented with 10 ng/mL IL-2 for 1h at 37°C, 90% humidity and 5% CO_2_. Then, cells were washed with PBS and stained for the respective markers using 1 μL of each antibody in 50 μL FACS buffer. Antibodies and isotype controls were obtained from BioLegend: Brilliant Violet 421™ anti-human CXCR4 (#306517), APC anti-human CD69 Antibody (#310910), PE anti-human CD25 Antibody (#302606), FITC anti-human CCR5 Antibody (#313705), APC anti-human GPR15 Antibody (#373006), Brilliant Violet 421™ anti-human CXCR6 (#356014). The recommended isotype control of each antibody described above was used at equal concentrations. Cells were incubated with the antibodies for 1 h at 4°C. Cells were washed and fixed in 4% (v/v) PFA in PBS. PBMCs were gated based on forward and side scatter characteristics, followed by exclusion of doublets and then by the respective marker expression. Data were generated with BD FACS Diva 6.1.3 Software before analysis with FlowJo 8.8 Software (Treestar).

#### Proteolysis of Cystatin C

100 μg (7.54 nmol) human Cystatin C protein (purified, from human urine, BML-SE479-0100) obtained from Enzo Life Sciences were digested with indicated proteases at 1:100 molar ratio (75.4 pmol protease). Digestions were performed in 0.2 M citrate buffer pH 4.0 (cathepsin D, G and E); 100 mM Tris HCl with 10 mM CaCl2 pH 8 (trypsin), 20 mM sodium acetate pH 3.5 (pepsin), 50 mM Tris HCl with 0.26 M NaCl pH 8 (chymase) or 0.1 M NaOAc with 0.2 M NaCl pH 3.5 (napsin A). Digestions with Cathepsin G and E were incubated for 24 h at 37°C, all other reactions for 2 h at 37°C. To remove proteases and exchange buffer, digestion products were diluted in PBS (75 μl digestions reaction mixture + 425 μl PBS) and applied to 30 kDa Amicon ultrafiltration units (Merck, #UFC5030). Samples were centrifuged for 30 minutes at 12,000 x g and room temperature. Columns were discarded and flow through 3 kDa Amicon ultrafiltration units (Merck, #UCF5003) was applied to retain CysC fragments (~ 5-7 kDa) and to remove salts and buffer components. Samples were centrifuged again as described above. Cystatin C peptides were eluated from the filter by inverting the unit in fresh 1.5 ml Eppendorf tubes. Samples were centrifuged for 2 minutes at 1,000 × g and eluate volumes adjusted to the starting volume of 75 μl. The amount of recovered digestion product used for the GHOST Bob infection assay was based on the molecular weight of the full-length Cystatin C and assuming complete digestion of the protein.

#### Coomassie staining

Digestion products were mixed with Protein Loading Buffer (LI-COR #928-40004) and TCEP (50 mM final concentration) and heated to 70°C for 10 min. Proteins were then separated on NuPAGE 4-12% BisTris gels, fixed with 50% methanol plus 7% acetic acid solution and stained with GelCode Blue (Thermo Fisher #24590) for 1 h at RT. After destaining with ultrapure water, the gel was imaged on the Gel Doc™ XR+ Gel Documentation System (BioRad).

#### GPR15 expression constructs

For all PCRs, the Phire Hot Start II DNA Polymerase (Thermo Fisher Scientific, F122L) was used. GPR15 was PCR-amplified from the plasmid pCMV6-XL4 human GPR15 (OriGene, CAT# SC116846, ACCN: NM-005290). Thereby the single restriction sites NheI and HindIII were added to the sequences by the primers used (Primer: NheI-GPR15-for and GPR15-HindIII-rev). The amplified GPR15 was cloned in the empty pcDNA3.1 vector by using the introduced single restriction sites. Additionally, GPR15 constructs C-or N-terminal tagged with eGFP were cloned via overlap-extension PCR. For the N-terminal tagged pcDNA3.1-eGFP-GPR15 construct eGFP was fused to GPR15 by replacing the stop codon of eGFP and the start codon of GPR15 by the linker sequence CCCGGG (Primer: eGFP NheI-for, GPR15-HindIII-rev, GFP-GPR15-SOE-rev and GFP-GPR15-SOE-for) and afterwards ligated in the pcDNA3.1 construct by using NheI and HindIII. The C-terminal tagged construct pcDNA3.1-GPR15-eGFP was constructed in the same way by fusing eGFP to the C-terminal end of GPR15 by replacing the stop codon of GPR15 and the start codon of eGFP with the linker sequence GGCGCCGGCGCC (Primer: NheI-GPR15-for, eGFP-HindIII-rev, GPR15-GFP-SOE-rev and GPR15-GFP-SOE-for).

#### CCR5 genotyping

1×10^7^ Δ32/Δ32 PBMCs were transferred into a 1.5 mL Eppendorf tube and spinned down. Supernatants were discarded and cells were washed three times in 1x PBS. The cell pellet was lysed in M-PER Mammalian Protein Extraction reagent (Thermo Fisher # 78501) in presence of 100 μg/mL Proteinase K for 1 h at 56°C, followed by an inactivation step at 95°C for 10 minutes. Polymerase chain reaction (PCR) was carried out using the Phusion High-Fidelity DNA Polymerase according to the manufacturers’ instructions using the primer pair p5CCR5fw 5’-TGG-TGG-CTG-TGT-TTG-CGT-CTC-3’ and p3CCR5rev 5’-AGC-GGC-AGG-ACC-AGC-CCC-AAG-3’. A total of 35 thermal cycles per PCR were performed in a VeritiTM 96-Well Thermal Cycler (Applied Biosystems) in a total volume of 50 μl. Then, 25 μl of product were separated on a 3 % agarose gel (w/v, Sigma Aldrich) in 1x TAE buffer (Carl Roth) next to a 1 kb Plus DNA ladder (Thermo Scientific). The samples were run for 25 min at 140 V, stained using ethidium bromide (AppliChem GmbH) and visualized in a Bio-Rad Gel Doc XR+

#### Confocal microscopy

5,000 Hela cells were seeded in 300 μL DMEM supplemented with FCS (10%), L-glutamine (2 mM), streptomycin (100 mg/mL) and penicillin (100 U/mL) in μ-Slide 8-well Ibidi microscopy chambers one day prior to the experiment. Cells were washed in PBS and medium was replaced with fresh medium once. Cells were then transfected with 0.25 μg GFP-tagged GPR15 expression plasmids using the TransIT®-LT1 Transfection Reagent (Mirus) according to the manufacturer’s protocol. 4 hours post-transfection, the medium was changed and cells were cultured for 2 days at 37°C, 90% humidity and 5% CO_2_. At 2 days post-transfection, cells were washed in PBS, fixed in 4% (v/v) PFA for 30 minutes at room temperature, permeabilized using Block and Perm Buffer (5% (v/v) FCS in PBS with 0.5% (v/v) Triton X100). Nuclei were stained using DAPI by incubating cells for 2 hours at 4°C. Cells were washed three times with PBS and once with HPLC water. Cells were covered with glycerol-based mounting medium to prevent bleaching of the fluorophores during microscopy. Confocal microscopy was performed using a LSM710 (Carl Zeiss). To examine the interaction of CysC95-146 with GPR15, Hela cells were transfected as described above. 2 days post-transfection, cells were treated with 5 μM Atto647-labeled CysC95-146 for 1 h at 4°C, then washed several times in ice-cold PBS, fixed in 4% (v/v) PFA for 30 minutes at 4°C. Nuclei were stained using

Hoechst33342 (Merck) according to the manufacturer’s instructions. Cells were washed with PBS, covered with glycerol-based mounting medium to prevent bleaching of the fluorophores during microscopy. Confocal microscopy was performed using an LSM710 (Carl Zeiss). Images were deconvoluted using Huygens Software (Scientific Volume Imaging) and processed in Fiji (Fiji is just ImageJ).

#### Fluorescence-based calcium release assays

GPR15 signaling efficiency was determined as described previously (Suply et al., 2017).

#### Antibody-mediated inhibition of GPR15-dependent virus infection

Antibodies targeting the N-terminus (ab8104) or the first extracellular loop (ab188938) were obtained from Abcam. An additional antibody against human GPR15 with unknown target site from R&D was also purchased (R&D #367902). GHOST-GPR15 cells were seeded 24h earlier at a density of 6,000 cells/well in 96 well F-bottom plates. Cells were preincubated with increasing amounts of anti-GPR15 antibodies for 2h at 37°C. Then, cells were infected with SIVmac239 F-Luc. At 3 dpi, infectivity was analyzed via firefly luciferase reporter assay.

#### CXCR4 Antibody competition assay

Competition of compounds with antibody binding was performed on SupT1 cells. Cells were washed in PBS containing 1 % FCS and 50,000 cells were then seeded per well in a 96 V-well plate. Buffer was removed and plates were precooled at 4°C. Compounds were diluted in PBS and antibody (clone 12G5, APC labeled) was diluted in PBS containing 1 % FCS. The antibody was used at a concentration close to its determined equilibrium dissociation constant (Kd). Compounds were added to the cells and 15 μl antibody immediately afterwards. Plates were incubated at 4°C in the dark for 2 hrs. Next, cells were washed with PBS containing 1% FCS and fixed with 2% PFA. Antibody binding was analyzed by flow cytometry (CytoFLEX; Beckman Coulter®).

#### Molecular Modeling

Based on the full CysC structure (Protein Data Bank identification code 3GAX), taken from the Protein Data Bank (Bernstein et al., 1977), the initial atomic positions of the CysC95-146 model were obtained. Possible initial GPR15 atomic coordinates were created with Phyre2 (Kelley et al., 2015), while hydrogen termination was performed with Maestro (www.schrodinger.com/releases/release-2019-3) for pH 7. Initial equilibration (300K for 0.1 ns) was then performed by classical MD (molecular dynamics) simulations using the GROMACS MD engine and the non-reactive force field Amber99sb. Afterwards, ReaxFF (reactive molecular dynamic) simulations (van Duin et al., 2001) were performed with the Amsterdam Modeling Suite 2019 (ADF2019, SCM, Theoretical Chemistry, Vrije Universiteit, Amsterdam, The Netherlands, http://www.scm.com) within *NVT* ensembles over 25 ps and coupling to a Berendsen heat bath (*T*=300 K with a coupling constant of 100 fs). Random interaction structures between GPR15 and CysC95-146 were generated and preselected using the overall system energy as selection criterion. For the most stable systems, reactive MD simulations were performed to evaluate the interaction energy and morphology. Afterwards, amino acid-resolved interaction energies were obtained by individually removing amino acids from the frustrated interaction model, followed by evaluating the energy changes. For all visualizations the Visual Molecular Dynamics program (VMD) (Humphrey et al., 1996) was used.

#### Statistical methods

The mean activities were compared using Student’s t-test. Similar results were obtained with the Mann Whitney test. The software package StatView version 4.0 (Abacus Concepts, Berkeley, CA, USA) was used for all calculations.

### QUANTIFICATION AND STATISTICAL ANALYSIS

Statistical analyses were performed with GraphPad PRISM (GraphPad Software) and Microsoft Excel. P values were calculated using the two-tailed unpaired Student’s-t-test or Mann Whitney test. Correlations were calculated with the linear regression module. Unless otherwise stated, all experiments were performed three times and data are shown as mean ± SEM. Significant differences are indicated as: *p < 0.05; **p < 0.01; ***p < 0.001. Statistical parameters are specified in the figure legends.

## KEY RESOURCE TABLE

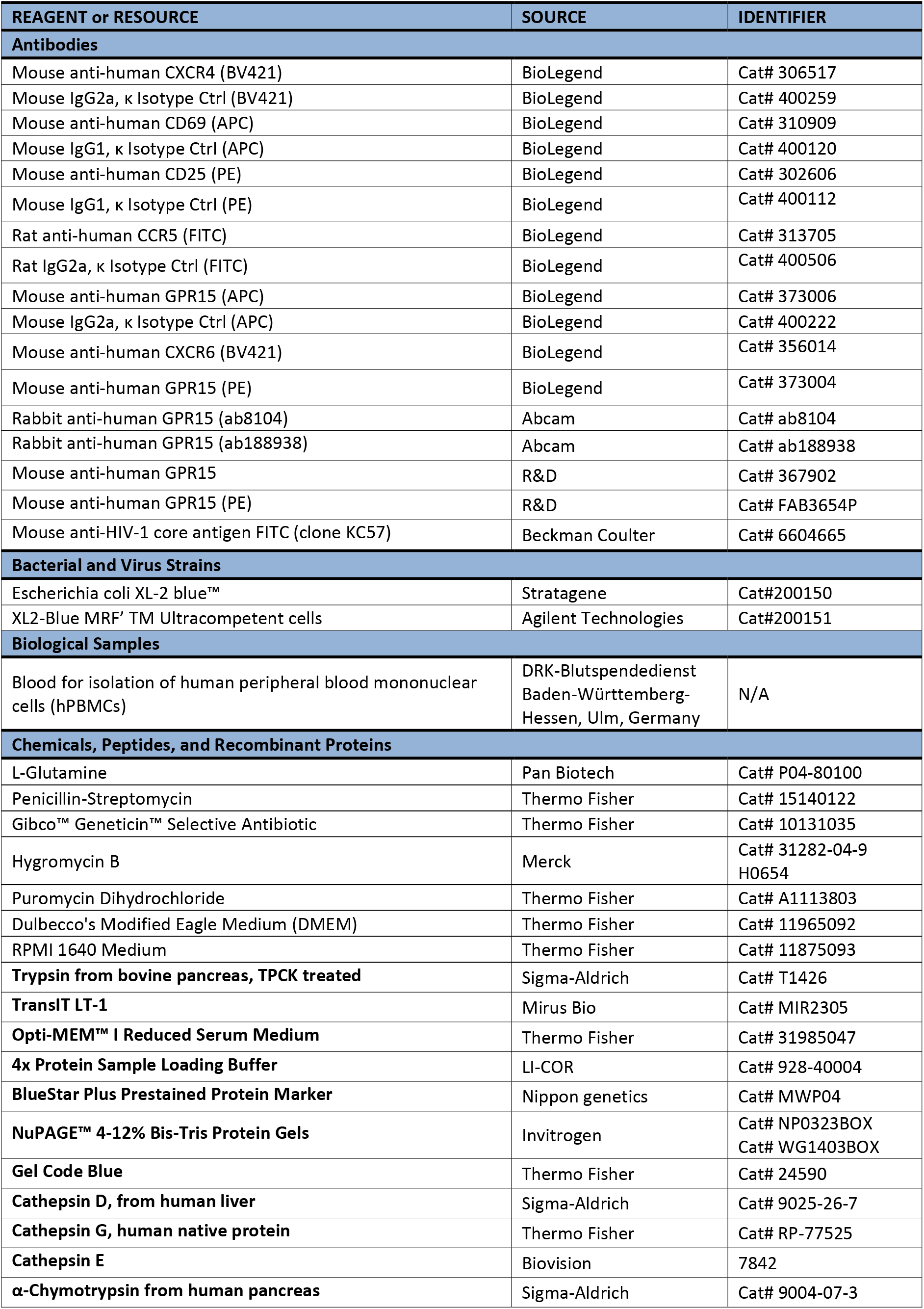
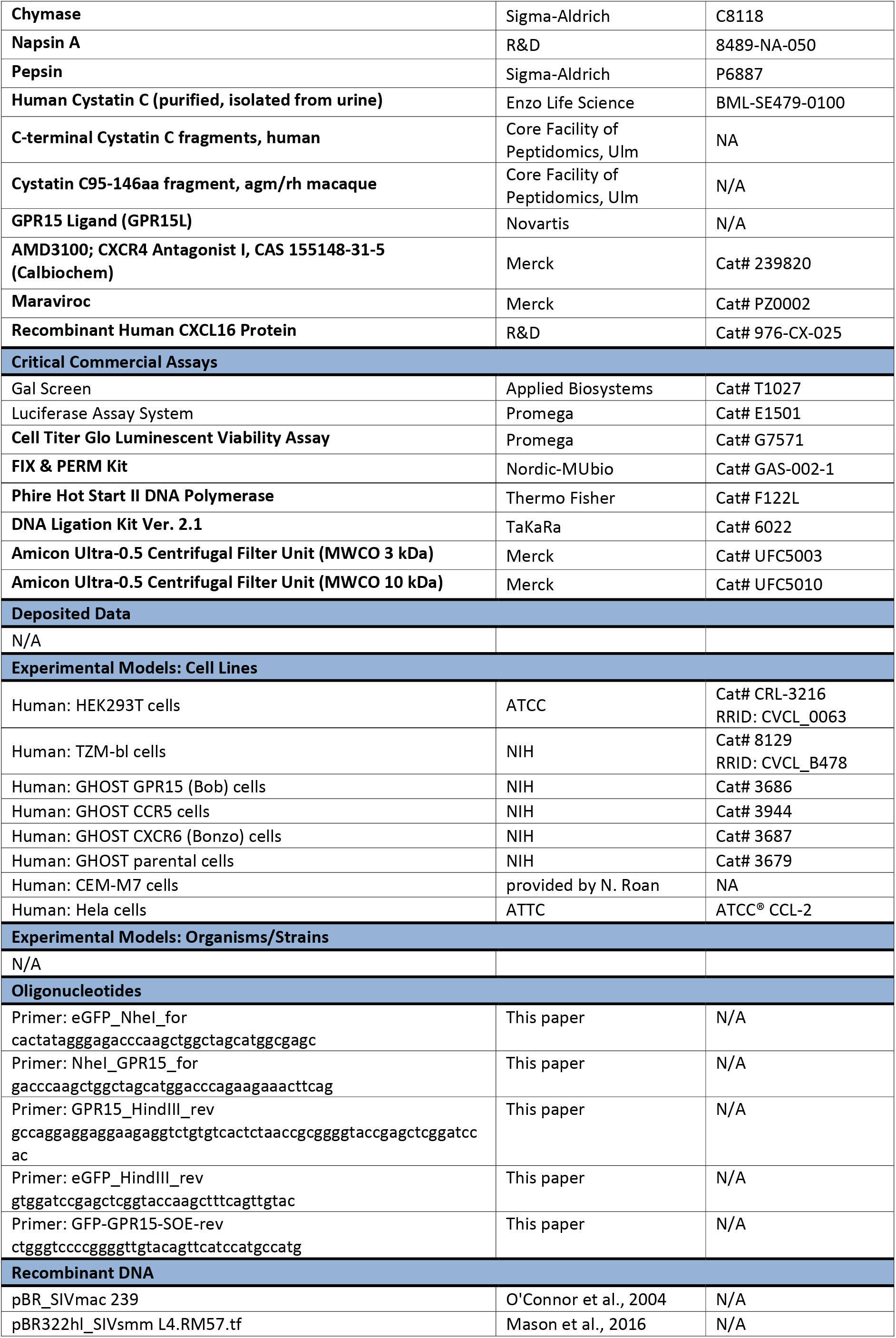
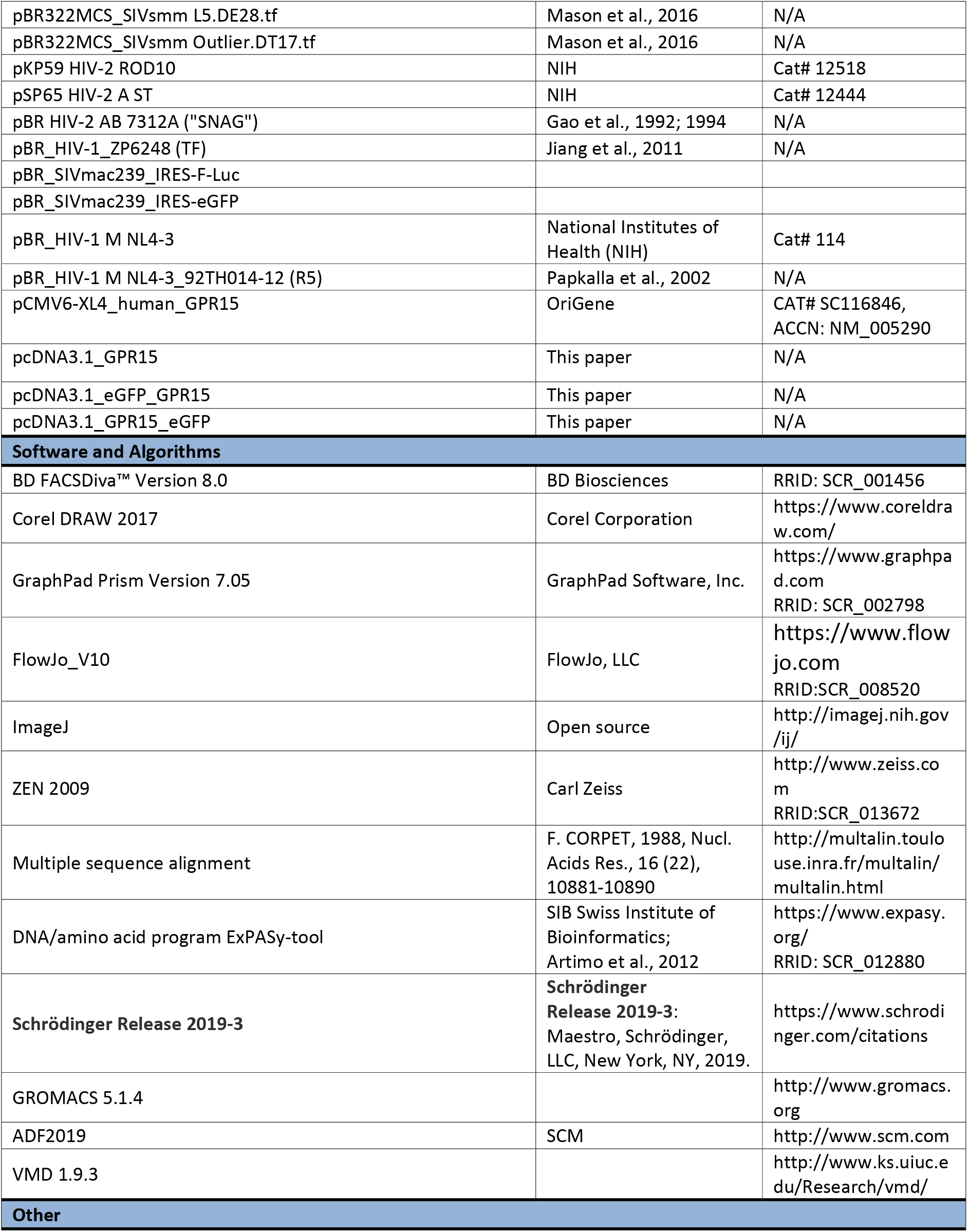

## Supplemental Figures

**Figure S1.**
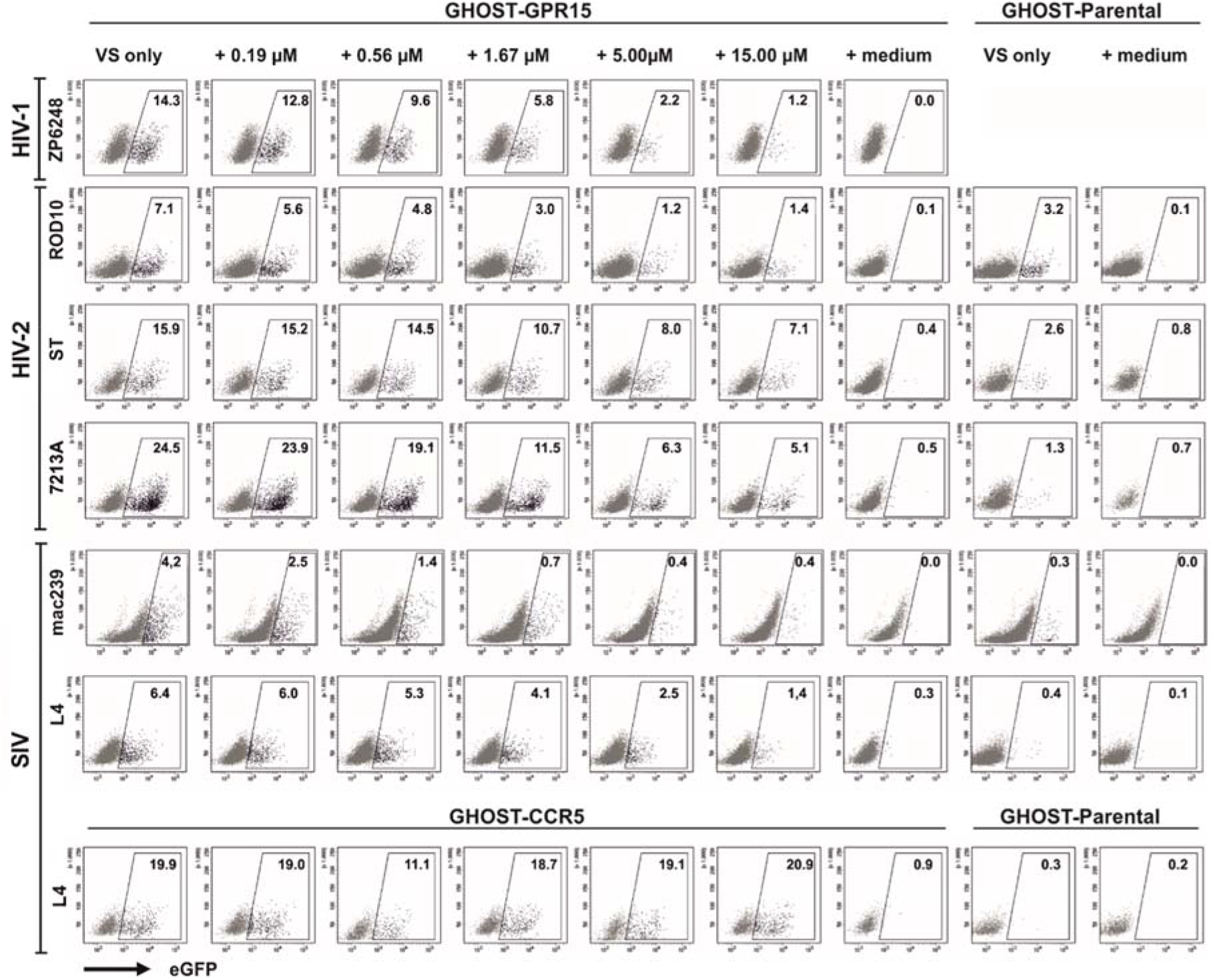
(related to Figure 2). Examples of primary FACS data for infection of GHOST cells in the presence of CysC95-146. GHOST-GPR15 or parental cells were infected with the indicated virus in the presence or absence of CysC95-146. Three days post-infection, infection rates were determined by flow cytometry. Exemplary primary data show the frequency of detected GFP-positive (infected) cells of the respective population.

**Figure S2.**
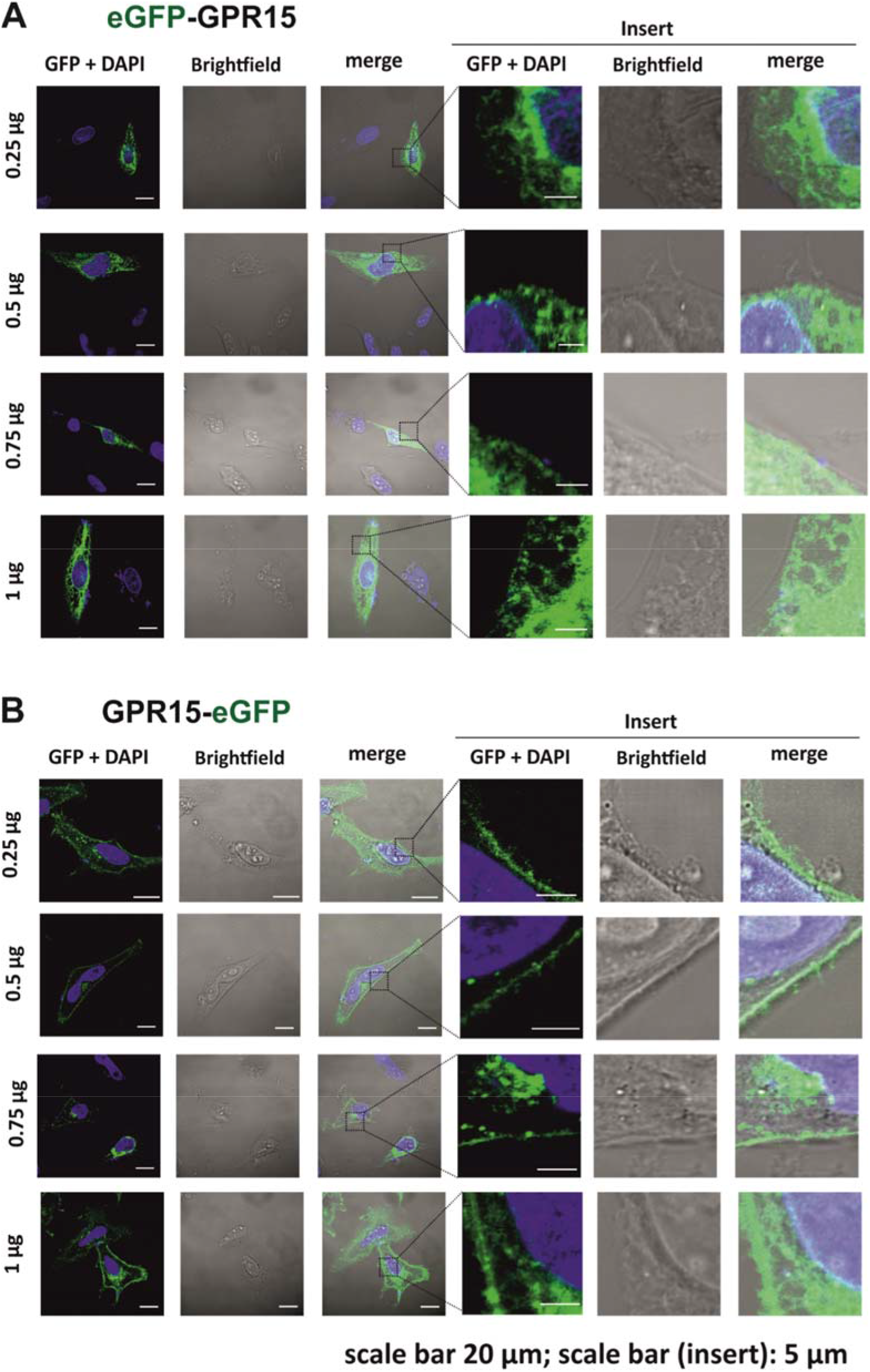
(related to Figure 3). Expression of eGFP-tagged GPR15. (**A, B**) Hela cells were transfected with the indicated amount of plasmid DNA of constructs expressing (A) N-terminally or (B) C-terminally eGFP-tagged GPR15. 2 days post-transfection, nuclei were stained using DAPI and analysed by confocal microscopy using an LSM710 (Carl Zeiss). Scale bar indicates 20 μm in the left panel and 5 μm in the insert panel.

**Figure S3.**
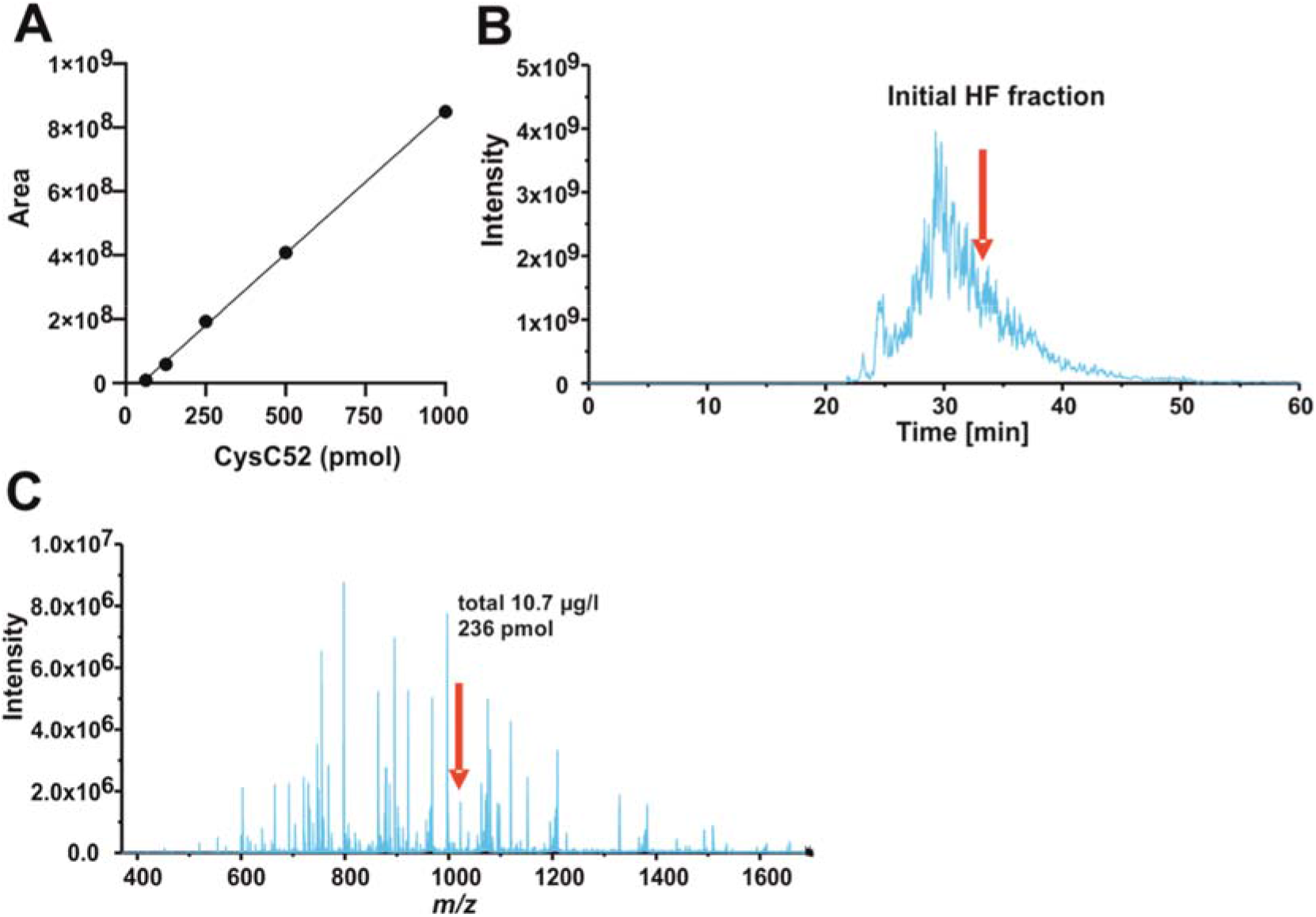
(related to Figure 5). Detection of CysC95-146 in the HF peptide library. (**A**) In “X” color: total ion chromatogram (TIC) of the starting HF sample (HF020802 P4 Fr19-20) visualized by XCalibur2.2 from the raw data obtained in the nanoLC-ESI-iTrap-Orbitrap system, and in “Y” color: ion chromatogram obtained after filtering TIC (m/z range 1025.3112±20 ppm; z=6) aiming to illustrate the CysC-F52 position in the starting HF chromatogram. (**B**) Mass spectrum of the components present within the time range where CysC95-146 is located (signal highlighted). (**C**) Calibration curve (peak area vs CysC95-146 amount) obtained by the analysis of CysC95-146 standards and the starting material HF 020802 P4Fr19-20, for the quantitation of CysC95-146 in hemofiltrate.

**Figure S4.**
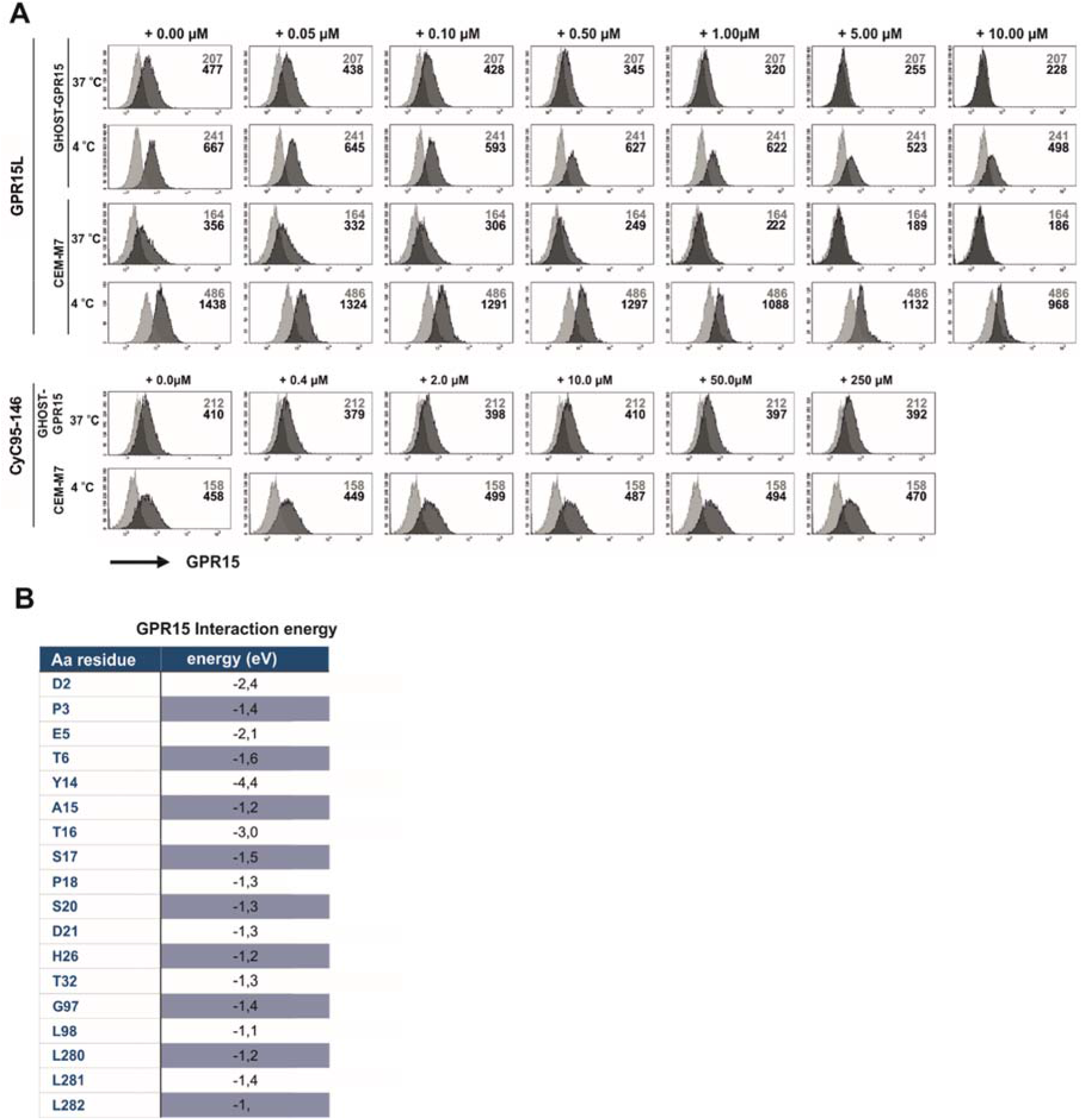
(related to Figure 6). Effect of GPR15L and CysC95-146 on GPR15 expression. (**A**) Changes in GPR15 surface expression on GHOST-GPR15 and CEM-M7 cells in the presence of GPR15L (Novartis) or CysC95-146. GHOST-GPR15 or CEM-M7 cells were incubated with different concentrations of GPR15L or CysC95-146 for 30 min at either 37°C or 4°C (to prevent receptor internalization) prior to staining with anti-GPR15 or isotype control antibody. GPR15 expression was analyzed by flow cytometry. Shown are the histograms for GPR15 (dark gray) and the respective isotype control (light gray). Numbers indicate the MFIs. (**B**) Predicted interaction energies of specific amino acid residues in GPR15 with CysC95-146.

**Figure S5.**
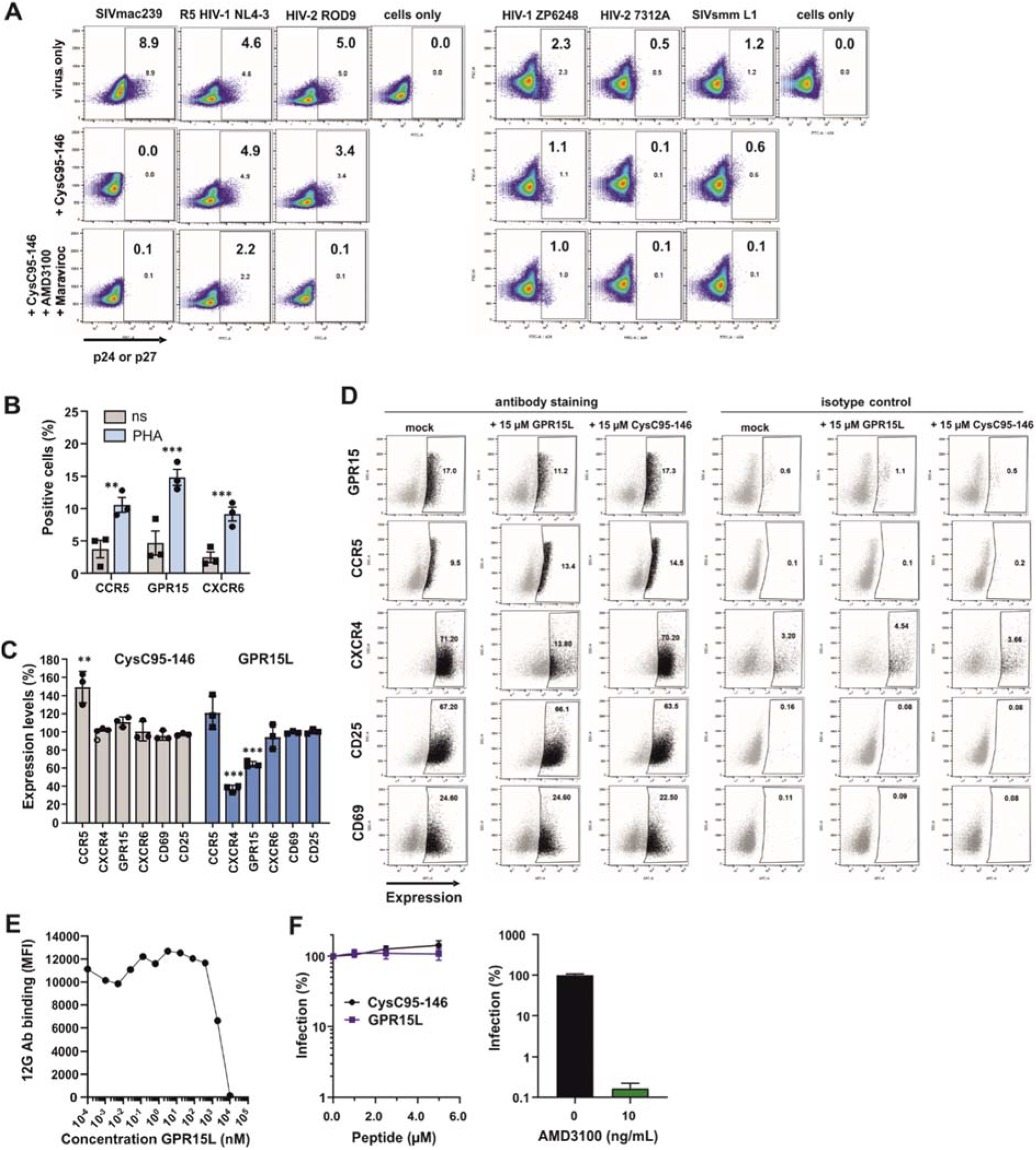
(related to Figure 7). Effect of CysC95-146 and GPR15L on virus infection and receptor expression in PBMC cultures. (**A**) CysC95-146 inhibits GPR15-mediated SIVmac239 infection in human PBMCs. Peripheral blood mononuclear cells (PBMCs) were isolated from buffy coats of three healthy donors, stimulated with PHA and IL-2. Cells were incubated with the indicated amounts of CysC95-146 or a combination of CysC95-146, AMD3100 and Maraviroc for 2h before infection with the indicated viruses. 3 dpi, PBMCs were stained for p24 and analyzed by flow cytometry. Shown are exemplary FACS data for one donor. Indicated with numbers are the frequency of infected cells. **(B)** Upregulation of CCR5, GPR15 and CXCR6 in human PBMCs upon stimulation with IL-2/PHA. PBMCs from three donors were isolated from buffy coats as described previously. Cells were then either analyzed for CCR5, GPR15 and CXCR6 expression immediately after the isolation or after a 3 days stimulation with IL-2/PHA. Shown are the mean values of n=3 donors. **(C)** Changes in the expression levels of different chemokine receptors and/or activation markers in the presence of GPR15L or CysC95-146. Data show normalized expression levels detected in stimulated PBMCs relative to the absence of peptide treatment (100%) of n = 3 donors +/- SEM, *** p < 0.001, ** p < 0.01, * p < 0.05 (Multiple t-test, unpaired, corrected for multiple comparisons using the Holm-Sidak method). (**D**) Exemplary FACS data of one donor showing the changes in surface expression of GPR15, CCR5, CXCR6, CD25, CD69 and CXCR4 in the presence of CysC95-146 or GPR15L as described in (C). **(E)** Competition of GPR15L with binding of Ab 12G5 targeting the 2^nd^ extracellular loop of CXCR4. (**F**) Effect of CysC95-146 and GPR15L (left) or AMD3100 (right) on CXCR4-tropic HIV-1 NL4-3 infection of GHOST-CXCR4 cells. Results were derived from three experiments and show mean values (±SEM) compared to the infection rates measured in the absence of inhibitor (100%).

## REFERENCES

Adri C. T. van Duin, †,ǁ, Siddharth Dasgupta, ‡, Francois Lorant, § and, and William A. Goddard III*, ‡ (2001). ReaxFF: A Reactive Force Field for Hydrocarbons.

Alkhatib, G., Combadiere, C., Broder, C.C., Feng, Y., Kennedy, P.E., Murphy, P.M., and Berger, E.A. (1996). CC CKR5: A RANTES, MIP-1, MIP-1 Receptor as a Fusion Cofactor for Macrophage-Tropic HIV-1. Science (80-.). 272, 1955–1958.

Appelqvist, H., Wäster, P., Kågedal, K., and Öllinger, K. (2013). The lysosome: from waste bag to potential therapeutic target. J. Mol. Cell Biol. 5, 214–226.

Bernstein, F.C., Koetzle, T.F., Williams, G.J., Meyer, E.F., Brice, M.D., Rodgers, J.R., Kennard, O., Shimanouchi, T., and Tasumi, M. (1977). The Protein Data Bank: a computer-based archival file for macromolecular structures. J. Mol. Biol. 112, 535–542.

Bhasin, B., Lau, B., Atta, M.G., Fine, D.M., Estrella, M.M., Schwartz, G.J., and Lucas, G.M. (2013). HIV Viremia and T-Cell Activation Differentially Affect the Performance of Glomerular Filtration Rate Equations Based on Creatinine and Cystatin C. PLoS One 8, e82028.

Bishop, J.A., Sharma, R., and Illei, P.B. (2010). Napsin A and thyroid transcription factor-1 expression in carcinomas of the lung, breast, pancreas, colon, kidney, thyroid, and malignant mesothelioma. Hum. Pathol. 41, 20–25.

Bosso, M., Ständker, L., Kirchhoff, F., and Münch, J. (2017). Exploiting the human peptidome for novel antimicrobial and anticancer agents. Bioorg. Med. Chem.

Chahroudi, A., Bosinger, S.E., Vanderford, T.H., Paiardini, M., and Silvestri, G. (2012). Natural SIV Hosts: Showing AIDS the Door. Science (80-.). 335, 1188–1193.

Choe, H., Farzan, M., Sun, Y., Sullivan, N., Rollins, B., Ponath, P.D., Wu, L., Mackay, C.R., LaRosa, G., Newman, W., et al. (1996). The beta-chemokine receptors CCR3 and CCR5 facilitate infection by primary HIV-1 isolates. Cell 85, 1135–1148.

Clavel, F., Guyader, M., Guétard, D., Sallé, M., Montagnier, L., and Alizon, M. (1986). Molecular cloning and polymorphism of the human immune deficiency virus type 2. Nature 324, 691–695.

Compton, A.A., Malik, H.S., and Emerman, M. (2013). Host gene evolution traces the evolutionary history of ancient primate lentiviruses. Philos. Trans. R. Soc. B Biol. Sci. 368, 20120496–20120496.

Connor, R.I., Sheridan, K.E., Ceradini, D., Choe, S., and Landau, N.R. (1997). Change in coreceptor use correlates with disease progression in HIV-1--infected individuals. J. Exp. Med. 185, 621–628.

Deng, H., Liu, R., Ellmeier, W., Choe, S., Unutmaz, D., Burkhart, M., Di Marzio, P., Marmon, S., Sutton, R.E., Hill, C.M., et al. (1996). Identification of a major co-receptor for primary isolates of HIV-1. Nature 381, 661–666.

Deng, H.K., Unutmaz, D., KewalRamani, V.N., and Littman, D.R. (1997). Expression cloning of new receptors used by simian and human immunodeficiency viruses. Nature 388, 296–300.

Detheux, M., Ständker, L., Vakili, J., Münch, J., Forssmann, U., Adermann, K., Pöhlmann, S., Vassart, G., Kirchhoff, F., Parmentier, M., et al. (2000). Natural proteolytic processing of hemofiltrate CC chemokine 1 generates a potent CC chemokine receptor (CCR)1 and CCR5 agonist with anti-HIV properties. J. Exp. Med. 192, 1501–1508.

Donzella, G.A., Schols, D., Lin, S.W., Esté, J.A., Nagashima, K.A., Maddon, P.J., Allaway, G.P., Sakmar, T.P., Henson, G., De Clercq, E., et al. (1998). AMD3100, a small molecule inhibitor of HIV-1 entry via the CXCR4 co-receptor. Nat. Med. 4, 72–77.

Doranz, B.J., Rucker, J., Yi, Y., Smyth, R.J., Samson, M., Peiper, S.C., Parmentier, M., Collman, R.G., and Doms, R.W. (1996). A dual-tropic primary HIV-1 isolate that uses fusin and the beta-chemokine receptors CKR-5, CKR-3, and CKR-2b as fusion cofactors. Cell 85, 1149–1158.

Döring, M., Borrego, P., Büch, J., Martins, A., Friedrich, G., Camacho, R.J., Eberle, J., Kaiser, R., Lengauer, T., Taveira, N., et al. (2016). A genotypic method for determining HIV-2 coreceptor usage enables epidemiological studies and clinical decision support. Retrovirology 13, 85.

Dorr, P., Westby, M., Dobbs, S., Griffin, P., Irvine, B., Macartney, M., Mori, J., Rickett, G., Smith-Burchnell, C., Napier, C., et al. (2005). Maraviroc (UK-427,857), a Potent, Orally Bioavailable, and Selective Small-Molecule Inhibitor of Chemokine Receptor CCR5 with Broad-Spectrum Anti-Human Immunodeficiency Virus Type 1 Activity. Antimicrob. Agents Chemother. 49, 4721–4732.

Dragic, T., Litwin, V., Allaway, G.P., Martin, S.R., Huang, Y., Nagashima, K.A., Cayanan, C., Maddon, P.J., Koup, R.A., Moore, J.P., et al. (1996). HIV-1 entry into CD4+ cells is mediated by the chemokine receptor CC-CKR-5. Nature 381, 667–673.

Farzan, M., Choe, H., Martin, K., Marcon, L., Hofmann, W., Karlsson, G., Sun, Y., Barrett, P., Marchand, N., Sullivan, N., et al. (1997). Two orphan seven-transmembrane segment receptors which are expressed in CD4-positive cells support simian immunodeficiency virus infection. J. Exp. Med. 186, 405–411.

Feng, Y., Broder, C.C., Kennedy, P.E., and Berger, E.A. (1996). HIV-1 entry cofactor: functional cDNA cloning of a seven-transmembrane, G protein-coupled receptor. Science 272, 872–877.

Forssmann, W.-G., The, Y.-H., Stoll, M., Adermann, K., Albrecht, U., Tillmann, H.-C., Barlos, K., Busmann, A., Canales-Mayordomo, A., Giménez-Gallego, G., et al. (2010). Short-term monotherapy in HIV-infected patients with a virus entry inhibitor against the gp41 fusion peptide. Sci. Transl. Med. 2, 63re3.

Gao, F., Yue, L., Robertson, D.L., Hill, S.C., Hui, H., Biggar, R.J., Neequaye, A.E., Whelan, T.M., Ho, D.D., and Shaw, G.M. (1994). Genetic diversity of human immunodeficiency virus type 2: evidence for distinct sequence subtypes with differences in virus biology. J. Virol. 68, 7433–7447.

Gifford, R.J., Katzourakis, A., Tristem, M., Pybus, O.G., Winters, M., and Shafer, R.W. (2008). A transitional endogenous lentivirus from the genome of a basal primate and implications for lentivirus evolution. Proc. Natl. Acad. Sci. U. S. A. 105, 20362–20367.

Golding, H., Khurana, S., Yarovinsky, F., King, L.R., Abdoulaeva, G., Antonsson, L., Owman, C., Platt, E.J., Kabat, D., Andersen, J.F., et al. (2005). CCR5 N-terminal region plays a critical role in HIV-1 inhibition by Toxoplasma gondii-derived cyclophilin-18. J. Biol. Chem. 280, 29570–29577.

Harms, M., Habib, M.M., Nemska, S., Nicolò, A., Gilg, A., Preising, N., Sokkar, P., Carmignani, S., Raasholm, M., Weidinger, G., et al. (2020). An optimized derivative of an endogenous CXCR4 antagonist prevents atopic dermatitis and airway inflammation. BioRxiv 2020.08.28.272781.

Heng, B.C., Aubel, D., and Fussenegger, M. (2013). An overview of the diverse roles of G-protein coupled receptors (GPCRs) in the pathophysiology of various human diseases. Biotechnol. Adv. 31, 1676–1694.

Humphrey, W., Dalke, A., and Schulten, K. (1996). VMD: visual molecular dynamics. J. Mol. Graph. 14, 33–38, 27-28.

Ibe, S., Yokomaku, Y., Shiino, T., Tanaka, R., Hattori, J., Fujisaki, S., Iwatani, Y., Mamiya, N., Utsumi, M., Kato, S., et al. (2010). HIV-2 CRF01_AB: First Circulating Recombinant Form of HIV-2. JAIDS J. Acquir. Immune Defic. Syndr. 54, 241–247.

Jiang, C., Parrish, N.F., Wilen, C.B., Li, H., Chen, Y., Pavlicek, J.W., Berg, A., Lu, X., Song, H., Tilton, J.C., et al. (2011). Primary Infection by a Human Immunodeficiency Virus with Atypical Coreceptor Tropism. J. Virol. 85, 10669–10681.

Kelley, L.A., Mezulis, S., Yates, C.M., Wass, M.N., and Sternberg, M.J.E. (2015). The Phyre2 web portal for protein modeling, prediction and analysis. Nat. Protoc. 10, 845–858.

Kim, S. V, Xiang, W. V, Kwak, C., Yang, Y., Lin, X.W., Ota, M., Sarpel, U., Rifkin, D.B., Xu, R., and Littman, D.R. (2013). GPR15-mediated homing controls immune homeostasis in the large intestine mucosa. Science 340, 1456–1459.

Kong, L.I., Lee, S.W., Kappes, J.C., Parkin, J.S., Decker, D., Hoxie, J.A., Hahn, B.H., and Shaw, G.M. (1988). West African HIV-2-related human retrovirus with attenuated cytopathicity. Science 240, 1525–1529.

Krumbiegel, M., Kirchhoff, F., and P√∂hlmann, S. (1999). Coreceptor usage of BOB/GPR15 and Bonzo/STRL33 by primary isolates of human immunodeficiency virus type 1. J. Gen. Virol. 80, 1241–1251.

Kumar, P., Hui, H.X., Kappes, J.C., Haggarty, B.S., Hoxie, J.A., Arya, S.K., Shaw, G.M., and Hahn, B.H. (1990). Molecular characterization of an attenuated human immunodeficiency virus type 2 isolate. J. Virol. 64, 890–901.

Liu, R., Paxton, W.A., Choe, S., Ceradini, D., Martin, S.R., Horuk, R., MacDonald, M.E., Stuhlmann, H., Koup, R.A., and Landau, N.R. (1996). Homozygous defect in HIV-1 coreceptor accounts for resistance of some multiply-exposed individuals to HIV-1 infection. Cell 86, 367–377.

Lobritz, M.A., Ratcliff, A.N., Marozsan, A.J., Dudley, D.M., Tilton, J.C., and Arts, E.J. (2013). Multifaceted mechanisms of HIV inhibition and resistance to CCR5 inhibitors PSC-RANTES and Maraviroc. Antimicrob. Agents Chemother. 57, 2640–2650.

Longenecker, C.T., Kitch, D., Sax, P.E., Daar, E.S., Tierney, C., Gupta, S.K., and McComsey, G.A. (2015). Reductions in Plasma Cystatin C After Initiation of Antiretroviral Therapy Are Associated With Reductions in Inflammation. JAIDS J. Acquir. Immune Defic. Syndr. 69, 168–177.

Magister, S., and Kos, J. (2013). Cystatins in immune system. J. Cancer 4, 45–56.

Mohr, K.B., Zirafi, O., Hennies, M., Wiese, S., Kirchhoff, F., and Münch, J. (2015). Sandwich enzyme-linked immunosorbent assay for the quantification of human serum albumin fragment 408-423 in bodily fluids. Anal. Biochem. 476, 29–35.

Mörner, A., Björndal, A., Albert, J., Kewalramani, V.N., Littman, D.R., Inoue, R., Thorstensson, R., Fenyö, E.M., and Björling, E. (1999). Primary human immunodeficiency virus type 2 (HIV-2) isolates, like HIV-1 isolates, frequently use CCR5 but show promiscuity in coreceptor usage. J. Virol. 73, 2343–2349.

Münch, J., Ständker, L., Pöhlmann, S., Baribaud, F., Papkalla, A., Rosorius, O., Stauber, R., Sass, G., Heveker, N., Adermann, K., et al. (2002). Hemofiltrate CC chemokine 1[9-74] causes effective internalization of CCR5 and is a potent inhibitor of R5-tropic human immunodeficiency virus type 1 strains in primary T cells and macrophages. Antimicrob. Agents Chemother. 46, 982–990.

Münch, J., Ständker, L., Adermann, K., Schulz, A., Schindler, M., Chinnadurai, R., Pöhlmann, S., Chaipan, C., Biet, T., Peters, T., et al. (2007). Discovery and optimization of a natural HIV-1 entry inhibitor targeting the gp41 fusion peptide. Cell 129, 263–275.

Münch, J., Ständker, L., Forssmann, W.-G., and Kirchhoff, F. (2014). Discovery of modulators of HIV-1 infection from the human peptidome. Nat. Rev. Microbiol. 12, 715–722.

Neuhaus, J., Jacobs, Jr, D.R., Baker, J.V., Calmy, A., Duprez, D., La Rosa, A., Kuller, L.H., Pett, S.L., Ristola, M., Ross, M.J., et al. (2010). Markers of Inflammation, Coagulation, and Renal Function Are Elevated in Adults with HIV Infection. J. Infect. Dis. 201, 1788–1795.

Nguyen, L.P., Pan, J., Dinh, T.T., Hadeiba, H., O’Hara, E., Ebtikar, A., Hertweck, A., Gökmen, M.R., Lord, G.M., Jenner, R.G., et al. (2014). Role and species-specific expression of colon T cell homing receptor GPR15 in colitis. Nat. Immunol. 16, 207–213.

Ocón, B., Pan, J., Dinh, T.T., Chen, W., Ballet, R., Bscheider, M., Habtezion, A., Tu, H., Zabel, B.A., and Butcher, E.C. (2017). A Mucosal and Cutaneous Chemokine Ligand for the Lymphocyte Chemoattractant Receptor GPR15. Front. Immunol. 8, 1111.

Odden, M.C., Scherzer, R., Bacchetti, P., Szczech, L.A., Sidney, S., Grunfeld, C., and Shlipak, M.G. (2007). Cystatin C level as a marker of kidney function in human immunodeficiency virus infection: the FRAM study. Arch. Intern. Med. 167, 2213–2219.

Okajima, F. (2013). Regulation of inflammation by extracellular acidification and proton-sensing GPCRs. Cell. Signal. 25, 2263–2271.

Owen, S.M., Ellenberger, D., Rayfield, M., Wiktor, S., Michel, P., Grieco, M.H., Gao, F., Hahn, B.H., and Lal, R.B. (1998). Genetically divergent strains of human immunodeficiency virus type 2 use multiple coreceptors for viral entry. J. Virol. 72, 5425–5432.

Pandrea, I., Sodora, D.L., Silvestri, G., and Apetrei, C. (2008). Into the wild: simian immunodeficiency virus (SIV) infection in natural hosts. Trends Immunol. 29, 419–428.

Pöhlmann, S., Stolte, N., Münch, J., Ten Haaft, P., Heeney, J.L., Stahl-Hennig, C., and Kirchhoff, F. (1999). Co-receptor Usage of BOB/GPR15 in Addition to CCR5 Has No Significant Effect on Replication of Simian Immunodeficiency Virus In Vivo. J. Infect. Dis. 180, 1494–1502.

Pöhlmann, S., Lee, B., Meister, S., Krumbiegel, M., Leslie, G., Doms, R.W., and Kirchhoff, F. (2000). Simian immunodeficiency virus utilizes human and sooty mangabey but not rhesus macaque STRL33 for efficient entry. J. Virol. 74, 5075–5082.

Rajamäki, K., Nordström, T., Nurmi, K., Åkerman, K.E.O., Kovanen, P.T., Öörni, K., and Eklund, K.K. (2013). Extracellular Acidosis Is a Novel Danger Signal Alerting Innate Immunity via the NLRP3 Inflammasome. J. Biol. Chem. 288, 13410–13419.

Randers, E., Kristensen, H., Erlandsen, E.., and Danielsen, S. (1998). Serum cystatin C as a marker of the renal function. Scand. J. Clin. Lab. Invest. 58, 585–592.

Richter, R., Schulz-Knappe, P., Schrader, M., Ständker, L., Jürgens, M., Tammen, H., and Forssmann, W.G. (1999). Composition of the peptide fraction in human blood plasma: database of circulating human peptides. J. Chromatogr. B. Biomed. Sci. Appl. 726, 25–35.

Riddick, N.E., Hermann, E.A., Loftin, L.M., Elliott, S.T., Wey, W.C., Cervasi, B., Taaffe, J., Engram, J.C., Li, B., Else, J.G., et al. (2010). A novel CCR5 mutation common in sooty mangabeys reveals SIVsmm infection of CCR5-null natural hosts and efficient alternative coreceptor use in vivo. PLoS Pathog. 6, e1001064.

Riddick, N.E., Wu, F., Matsuda, K., Whitted, S., Ourmanov, I., Goldstein, S., Goeken, R.M., Plishka, R.J., Buckler-White, A., Brenchley, J.M., et al. (2015). Simian Immunodeficiency Virus SIVagm Efficiently Utilizes Non-CCR5 Entry Pathways in African Green Monkey Lymphocytes: Potential Role for GPR15 and CXCR6 as Viral Coreceptors. J. Virol. 90, 2316–2331.

Rodríguez, A., Webster, P., Ortego, J., and Andrews, N.W. (1997). Lysosomes Behave as Ca ^2+^-regulated Exocytic Vesicles in Fibroblasts and Epithelial Cells. J. Cell Biol. 137, 93–104.

Samson, M., Libert, F., Doranz, B.J., Rucker, J., Liesnard, C., Farber, M., Saragosti, S., Lapoumeroulie, C., Cognaux, J., Forceille, C., et al. (1996). Resistance to HIV-1 infection in caucasian individuals bearing mutant alleles of the CCR-5 chemokine receptor gene. Nature 382, 722–726.

Sauter, D., and Kirchhoff, F. (2019). Key Viral Adaptations Preceding the AIDS Pandemic. Cell Host Microbe 25, 27–38.

Schulz-Knappe, P., Schrader, M., Ständker, L., Richter, R., Hess, R., Jürgens, M., and Forssmann, W.G. (1997). Peptide bank generated by large-scale preparation of circulating human peptides. J. Chromatogr. A 776, 125–132.

Sharp, P.M., and Hahn, B.H. (2011). Origins of HIV and the AIDS Pandemic. Cold Spring Harb. Perspect. Med. 1, a006841–a006841.

Steen, A., W. Schwartz, T., and M. Rosenkilde, M. (2009). Targeting CXCR4 in HIV Cell-Entry Inhibition. Mini-Reviews Med. Chem. 9, 1605–1621.

Sun, H., Lou, X., Shan, Q., Zhang, J., Zhu, X., Zhang, J., Wang, Y., Xie, Y., Xu, N., and Liu, S. (2013). Proteolytic Characteristics of Cathepsin D Related to the Recognition and Cleavage of Its Target Proteins. PLoS One 8, e65733.

Suply, T., Hannedouche, S., Carte, N., Li, J., Grosshans, B., Schaefer, M., Raad, L., Beck, V., Vidal, S., Hiou-Feige, A., et al. (2017). A natural ligand for the orphan receptor GPR15 modulates lymphocyte recruitment to epithelia. Sci. Signal. 10, eaal0180.

Tissera, H., Rathore, A.P.S., Leong, W.Y., Pike, B.L., Warkentien, T.E., Farouk, F.S., Syenina, A., Eong Ooi, E., Gubler, D.J., Wilder-Smith, A., et al. (2017). Chymase Level Is a Predictive Biomarker of Dengue Hemorrhagic Fever in Pediatric and Adult Patients. J. Infect. Dis. 216, 1112–1121.

Turk, V., Stoka, V., Vasiljeva, O., Renko, M., Sun, T., Turk, B., and Turk, D. (2012). Cysteine cathepsins: From structure, function and regulation to new frontiers. Biochim. Biophys. Acta - Proteins Proteomics 1824, 68–88.

Unutmaz, D., KewalRamani, V.N., and Littman, D.R. (1998). G protein-coupled receptors in HIV and SIV entry: New perspectives on lentivirus-host interactions and on the utility of animal models. Semin. Immunol. 10, 225–236.

Venkatakrishnan, A.J., Deupi, X., Lebon, G., Tate, C.G., Schertler, G.F., and Babu, M.M. (2013). Molecular signatures of G-protein-coupled receptors. Nature 494, 185–194.

Verani, A., and Lusso, P. (2002). Chemokines as natural HIV antagonists. Curr. Mol. Med. 2, 691–702.

Villa, P., Jiménez, M., Soriano, M.-C., Manzanares, J., and Casasnovas, P. (2005). Serum cystatin C concentration as a marker of acute renal dysfunction in critically ill patients. Crit. Care 9, R139.

Visseaux, B., Damond, F., Matheron, S., Descamps, D., and Charpentier, C. (2016). Hiv-2 molecular epidemiology. Infect. Genet. Evol. 46, 233–240.

Woolley, M.J., and Conner, A.C. (2017). Understanding the common themes and diverse roles of the second extracellular loop (ECL2) of the GPCR super-family. Mol. Cell. Endocrinol. 449, 3–11.

Xiao, L., Rudolph, D.L., Owen, S.M., Spira, T.J., and Lal, R.B. (1998). Adaptation to promiscuous usage of CC and CXC-chemokine coreceptors in vivo correlates with HIV-1 disease progression. AIDS 12, F137–43.

Yamamoto, K., Kawakubo, T., Yasukochi, A., and Tsukuba, T. (2012). Emerging roles of cathepsin E in host defense mechanisms. Biochim. Biophys. Acta - Proteins Proteomics 1824, 105–112.

Zaidi, N., and Kalbacher, H. (2008). Cathepsin E: A mini review. Biochem. Biophys. Res. Commun. 367, 517–522.

Zhou, N., Luo, Z., Luo, J., Liu, D., Hall, J.W., Pomerantz, R.J., and Huang, Z. (2001). Structural and functional characterization of human CXCR4 as a chemokine receptor and HIV-1 co-receptor by mutagenesis and molecular modeling studies. J. Biol. Chem. 276, 42826–42833.

Zi, M., and Xu, Y. (2018). Involvement of cystatin C in immunity and apoptosis. Immunol. Lett. 196, 80–90.

Zirafi, O., Kim, K.-A., Ständker, L., Mohr, K.B., Sauter, D., Heigele, A., Kluge, S.F., Wiercinska, E., Chudziak, D., Richter, R., et al. (2015). Discovery and Characterization of an Endogenous CXCR4 Antagonist. Cell Rep. 11, 737–747.

